# Correlations enhance the behavioral readout of neural population activity in association cortex

**DOI:** 10.1101/2020.04.03.024133

**Authors:** Martina Valente, Giuseppe Pica, Caroline A. Runyan, Ari S. Morcos, Christopher D. Harvey, Stefano Panzeri

## Abstract

The spatiotemporal structure of activity in populations of neurons is critical for accurate perception and behavior. Experimental and theoretical studies have focused on “noise” correlations – trial-to-trial covariations in neural activity for a given stimulus – as a key feature of population activity structure. Much work has shown that these correlations limit the stimulus information encoded by a population of neurons, leading to the widely-held prediction that correlations are detrimental for perceptual discrimination behaviors. However, this prediction relies on an untested assumption: that the neural mechanisms that read out sensory information to inform behavior depend only on a population’s total stimulus information independently of how correlations constrain this information across neurons or time. Here we make the critical advance of simultaneously studying how correlations affect both the encoding and the readout of sensory information. We analyzed calcium imaging data from mouse posterior parietal cortex during two perceptual discrimination tasks. Correlations limited the ability to encode stimulus information, but (seemingly paradoxically) correlations were higher when mice made correct choices than when they made errors. On a single-trial basis, a mouse’s behavioral choice depended not only on the stimulus information in the activity of the population as a whole, but unexpectedly also on the consistency of information across neurons and time. Because correlations increased information consistency, sensory information was more efficiently converted into a behavioral choice in the presence of correlations. Given this enhanced-by-consistency readout, we estimated that correlations produced a behavioral benefit that compensated or overcame their detrimental information-limiting effects. These results call for a re-evaluation of the role of correlated neural activity, and suggest that correlations in association cortex can benefit task performance even if they decrease sensory information.

## Introduction

The collective activity of a population of neurons, beyond properties of individual cells, is thought to be critical for perceptual discrimination behaviors^1,2^. A fundamental question is how functional interactions in a population impact both the encoding of sensory information and how this information is read out downstream to guide behavioral choices^3^. A commonly studied feature of coding in neural populations is noise correlations, that is correlated trial-to-trial variability during repeated presentations of the same stimulus^4,5^. These correlations can take the form of correlations between the spike rates of two individual cells, which we call across-neuron noise correlations. Similarly, neural population activity at a given time in response to a stimulus is often correlated with the activity of the same population at other times, which we refer to as across-time noise correlations.

The impact of across-neuron and across-time correlations has been long debated. Much experimental and theoretical work has proposed that correlations limit the information capacity of a neural population^6–9^. Because these correlations reflect trial-to-trial variability that is shared across neurons or time, the detrimental effect of variability on stimulus decoding cannot be eliminated by averaging the activity of neurons or time points. Noise correlations are therefore proposed to fundamentally constrain what neural networks can compute, and limit the accuracy by which subjects can judge differences between stimuli^6,7^. A reason to suspect that the effect of noise correlations may be more nuanced, however, comes from a separate stream of biophysical and theoretical studies. This line of work has shown that spatially and temporally correlated spiking can more strongly and more reliably propagate signals and drive responses in postsynaptic neural populations^10–14^. However, it remains poorly tested if and how enhanced signal propagation may have a beneficial impact on behavioral discrimination performance.

We reasoned that we could investigate how noise correlations shape behavioral performance in perceptual discrimination by studying at the same time not only how correlations impact the encoding of sensory information, as has been emphasized frequently, but critically also how they impact the reading out of this information by downstream neural circuits to guide behavioral choices.

### Correlations of PPC activity limit sensory coding during perceptual discrimination

To examine how noise correlations affect both stimulus coding at the population level and behavioral discrimination performance, we focused on the mouse posterior parietal cortex (PPC). PPC is thought to participate in transforming multisensory signals into behavioral outputs and is thus likely at the interface of encoding and reading out stimulus information. Furthermore, PPC is essential for perceptual discrimination tasks during virtual-navigation^15,16^, and its activity has been shown to contain stimulus information that relates to an animal’s choices^16–22^. It is thus a relevant area to study the impact of correlated neural activity on behavior.

We examined across-time and across-neuron correlations in PPC population activity using previously published calcium imaging datasets. To study across-time correlations, we used calcium imaging data from a sound localization task^17^ in which mice reported perceptual decisions about the location of an auditory stimulus by navigating through a visual virtual reality T-maze (Fig. 1a). As mice ran down the T-stem, a sound cue was played from one of eight possible locations in head-centered, real-world coordinates. Mice reported whether the sound originated from their left or right by turning in that direction at the T-intersection (78.0 ± 0.5% correct). During each session, the activity of ∼50 layer 2/3 neurons was imaged simultaneously. This task had the same stimulus category (sound location) presented throughout the trial and thus was well suited to probe across-time correlations in neural activity. To study across-neuron correlations, we used calcium imaging data from an evidence accumulation task^19^ in which ∼350 layer 2/3 neurons were imaged simultaneously per session, thus providing a large population size to test across-neuron correlations. In the evidence accumulation task, during virtual navigation, mice were presented with six temporally separate visual cues on the left or right walls of a T-maze (Fig. 1f). Mice reported which side had more cues by turning in that direction at the T-intersection (84.5 ± 1.6% correct). We categorized the visual stimuli as having more total left or right cues. For all analyses, we focused on the period toward the end of the T-stem before the mouse had reported its choice and after it had received sensory information; this is a window in which neural activity may carry sensory information that is used to inform the choice (Methods).

**Figure 1:**
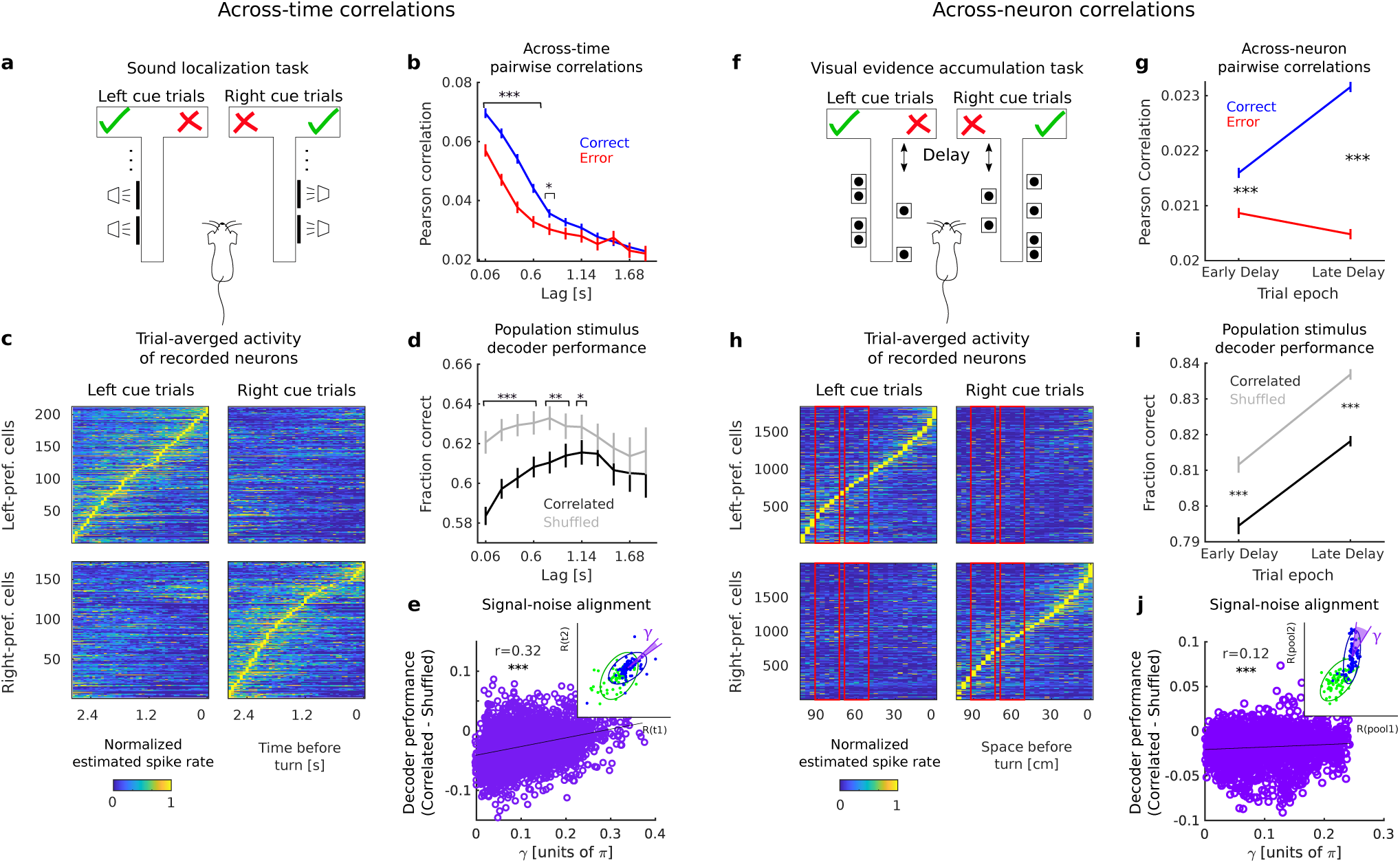
Across-time and across-neuron correlations in PPC responses limit sensory coding during perceptual discrimination tasks. **a**, Schematic of the sound localization task. Left/right sound category (speaker symbols) indicate the rewarded side of the maze (checkmark). **b**, Pearson’s correlations in time-lagged activity of all simultaneously-recorded pairs of PPC cells at fixed stimulus, computed separately for correct and error trials. Results are averaged across all time point pairs separated by a given lag. Data are mean ± s.e.m. across all imaged cell pairs. ***P<0.001; *P<0.05, two-sided permutation test. **c**, Normalized mean estimated spike rate traces for all the PPC cells imaged during the sound localization task, divided by left-preferring (n=212) and right preferring (n=172) cells, aligned to the turn frame. Traces were normalized to the peak of each cell’s mean estimated spike rate on preferred trials and sorted by peak time. **d**, Performance of a decoder of stimulus identity applied to joint population activity at two different time points, separately for real recorded (black) or trial-shuffled (gray) population vectors. Data are mean ± s.e.m. across sessions and time point pairs. ***P<0.001; **P<0.01; *P<0.05, two-sided permutation test. **e**, Correlation between difference in stimulus decoder performance from real and shuffled population vectors and misalignment angle *γ* between signal and noise. The inset shows an example of *γ* computation in the reduced 2D space of projected population activity vectors (see Methods). Ellipses indicate 90% confidence interval for best-fit bivariate Gaussian. ***P<0.001, circular-linear correlation^43^. **f**, Schematic of the evidence accumulation task. The rewarded side of the maze (checkmark) is the one identified by the most numerous visual cues (wall segments with black dots patterns). **g**, Pearson’s correlations in time-averaged activity of all simultaneously-recorded pairs of PPC cells, computed separately for correct and error trials. Data are mean ± s.e.m. across all imaged cell pairs. ***P<0.001, two-sided permutation test. **h**, As in **b**, for all cells (left-preferring, n=1840; right-preferring: n=2000) imaged during the evidence accumulation task. Note that activity traces are averaged over spatial bins. Red rectangles indicate the trial epochs selected for analyses (Early delay, Late delay). **i**, Performance of a decoder of stimulus identity applied to joint population activity of two randomly-defined non-overlapping neuronal pools, separately for real recorded (black) or within-pool trial-shuffled (gray) population vectors. Data are mean ± s.e.m. across sessions and 100 pairs of neuronal pools. ***P<0.001, two-sided permutation test. **j**, As in **e**. For panels **d-e, i-j**, data are first averaged across trial splits.

We tested how noise correlations impacted the encoding of sensory information in population activity. For the sound localization task, we quantified across-time correlations by computing the Pearson correlation between the activity of one neuron at one time point and the activity of another neuron at another time point, for each pair of neurons during trials with the same sound location category. Consistent with previous reports^17,19,23^, noise correlations were present even at lags longer than 1 s (Fig. 1b). Since many neurons exhibited activity selective for distinct trial types, with different neurons active at distinct time points in the trial (Fig. 1c), we were able to decode the stimulus category (left vs. right sound location) significantly above chance from pairs of temporally offset instantaneous population activity vectors (Methods; Fig. 1d). To evaluate how across-time correlations affected the encoding of stimulus category information, we shuffled instantaneous population activity vectors across trials of the same stimulus category, independently at each time point. This shuffle destroyed within-trial temporal relationships between population vectors while preserving instantaneous population activity (Methods). Importantly, stimulus decoding performance was higher when correlations were disrupted by shuffling, indicating that across-time correlations limited sound category information in neural activity (Fig. 1d). Similarly, for the evidence accumulation task, we computed across-neuron Pearson correlations between the activity of a pair of neurons in a given time window for trials with the same stimulus category (Fig. 1g). Because neurons exhibited activity selective to the stimulus category (more left or right cues) (Fig. 1h), we were able to decode the stimulus category from the population activity vector (Fig. 1i). We disrupted across-neuron correlations by dividing the neural population into two non-overlapping randomly-chosen pools of neurons of equal size and shuffling the trial labels separately for each pool within trials of the same stimulus category. This shuffle scrambled within-trial relationships between pools and thus removed correlations between pools (Methods). Importantly, decoding accuracy was higher when correlations were disrupted by shuffling than in the unshuffled data with correlations intact, showing that across-neuron correlations were also information limiting (Fig. 1i).

Whether noise correlations limit the information encoded by a neural population depends on how they relate to signal correlations (the correlations between trial-averaged responses to individual stimuli). We parametrized the relationship between signal and noise correlations in our data by computing the angle^24,25^, which we termed *γ*, between the axis of largest variation of neural responses at fixed stimulus (noise correlations axis) and the axis of largest stimulus-related variation (signal correlations axis) (Methods and Fig. 1e-j, inset). This angle was variable, but small on average (γ=0.111π ± 0.001π and 0.098π ± 0.001π for the sound localization and evidence accumulation datasets respectively, Fig. 1e, j). The smaller γ, the more noise correlations made the discrimination of different stimuli more difficult by increasing the overlap between the stimulus-specific response distributions to different stimuli (Fig. 1e, j, and Extended Data Fig. 2a, b). High γ values, corresponding to information-enhancing correlations reported in some previous studies^26,27^, were present also in our dataset, but were not predominant (Fig. 1e, j).

Often it has been assumed that perceptual discrimination performance increases proportionally with the amount of sensory information encoded in neural activity^28^. Accordingly, the fact that correlations limit sensory information has been interpreted to imply that correlations hinder the ability to discriminate sensory stimuli^6,29^. Specifically, across-time correlations have been proposed to limit the benefit of integrating noisy information over time as is commonly considered for a speed-accuracy tradeoff^30,31^. Further, across-neuron correlations are thought to lessen the benefit of averaging noisy information across populations of neurons^6,7,9^. If correlations are detrimental to perceptual behaviors, one would intuitively expect noise correlations to be lower when animals make correct choices and higher when animals make errors. Contrary to this expectation, both across-time and across-neuron correlations were higher in correct trials than in error trials (Fig. 1b, g), despite limiting information in population activity. This finding leads to the paradoxical suggestion that correlations limit information encoded by a neural population but at the same time may be beneficial for making accurate choices.

### A simple model of encoding and readout to test how correlations affect task performance

To reconcile the observations of correlations being information limiting, but at the same time being higher in correct trials, we developed a simple mathematical model that incorporated both the encoding of stimulus information and the readout of this information to form a choice. We compared two alternative views of how information in population activity may be used to perform a stimulus discrimination task. In the traditional view, the accuracy of a choice is proportional to the amount of information in a neural population, and thus information-limiting correlations constrain task performance. Alternatively, we considered that a choice could depend on both stimulus information and features of neural activity that emerge from correlations, in particular the consistency of information across time and neurons in a population.

We simulated a perceptual discrimination task with two possible stimuli that had to be converted into two possible corresponding choices (c = 1 for s = 1 and c = −1 for s = −1) (Fig. 2a). We simulated neural activity generically for two features r_1_ and r_2_, which could represent neural activity at different points in time (i.e. for across-time correlations) or activity of different neurons (i.e. for across-neuron correlations) (Fig. 2c). For each feature alone, higher-than-average values were simulated to indicate one sensory stimulus (s = 1), and lower-than-average values were simulated to indicate the opposite sensory stimulus (s = −1), meaning that the two features showed positive signal correlations. In addition, we simulated noise correlations between r_1_ and r_2_, such that the noise axis was closely aligned to the signal axis (γ = 0.08π, close to the mean values of both experimental datasets). Because of the close signal-noise alignment, noise correlations increased the overlap between the stimulus-specific response distributions (cf. blue and green ellipses in Fig. 2c, left panel vs right panel) and decreased the stimulus information encoded by the two features jointly (Fig. 2d).

**Figure 2:**
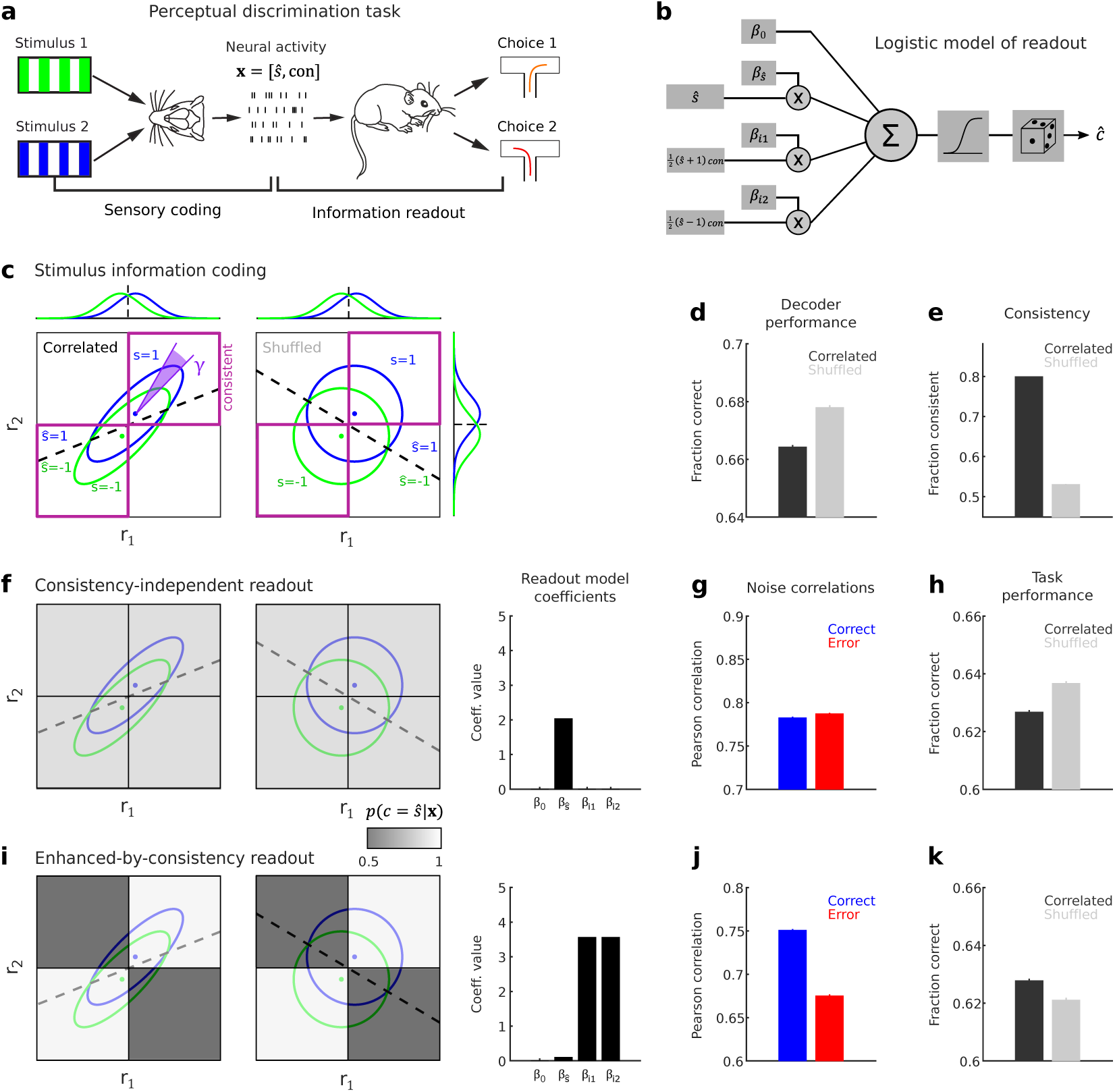
A simple encoding-readout model of stimulus information shows that different readouts determine different impacts of correlations on task performance. **a**, Schematic showing the two fundamental stages in the information processing chain for sensory perception, which are the constituting parts of the simple model that we propose: sensory coding and information readout. Sensory coding refers to the mapping from external stimuli (Stimulus 1, green; Stimulus 2, blue) to neural activity; information readout refers to the mapping from neural activity to a behavioral choice (Choice 1, turn right; Choice 2, turn left). Note that task-relevant neural activity is here recapitulated by two selected features, being the stimulus predicted when observing a given population activity (*ŝ*) and its consistency (*con*) (see Methods). **b**, Schematic of the readout model that was used to generate synthetic choices from the simulated neural activity patterns (see Methods). Task predictors extracted from simulated neural activity are weighted by corresponding model coefficients, summed and transformed non-linearly through a sigmoid function that outputs the probability of making a given binary choice. The actual choice is generated by randomly drawing from a binomial distribution with probability of single experiment success matching the output of the sigmoid function. **c**, Example of simulated response distributions along two neural features (r_1_, r_2_) to two stimuli (s=1, s=-1), modelled as bivariate Gaussians with different degree of correlation (correlated: ρ=.8, uncorrelated: ρ=0). For the correlated example, noise correlations axis and signal correlations axis are closely but not perfectly aligned (γ=0.08π). Ellipses indicate 95% confidence intervals. The dashed black line indicates the optimal stimulus decoding boundary. Purple squares indicate regions of the 2D response space in which r_1_ and r_2_ encode consistent stimulus information, i.e. they both code for the same stimulus (see Methods). Marginal response distributions along r_1_ and r_2_ and the corresponding decoding boundaries are shown on the sides. **d**, Fraction correct of a linear decoder of stimulus identity applied to simulated bivariate responses as in **c** (nr. trials = 1000, nr. simulations = 1000) is higher for uncorrelated responses (correlations limit the encoded stimulus information). **e**, The fraction of trials in which r_1_ and r_2_ encode consistent stimulus information is higher for correlated responses (correlations increase consistency). **f**, Consistency-independent readout represented as a grayscale map in the r_1_-r_2_ response space is superimposed to the distributions in **a**. The shade of gray represents the readout efficacy of transformation from encoded stimulus into choice (see Methods), which is the same for consistent and inconsistent trials (*p*(*c* = *ŝ*|*ŝ, con* = 1) = *p*(*c* = *ŝ*|*ŝ, con* = 0) = .89). The corresponding readout model coefficients are shown on the right. **g**, Pearson correlation coefficient of neural responses at fixed stimulus is slightly higher in trials with incorrect choice than in correct trials, when readout is consistency-independent and correlations are as in **c. h**, Task performance resulting from the consistency-independent readout of simulated neural activity is higher for uncorrelated responses, thus reflecting the impact of correlations on stimulus coding. **i**, Same as **f**, for enhanced-by-consistency readout (consistency modulation index *η* = 0.9, see Methods). The readout efficacy differs between consistent and inconsistent trials (*p*(*c* = *ŝ*|*ŝ, con* = 1) = .97; *p*(*c* = *ŝ*|*ŝ, con* = 0) = .53). **j**, Pearson correlation coefficient is higher in trials with correct choice than in error trials, when readout is enhanced-by-consistency and correlations are as in **c. k**, Task performance resulting from the enhanced-by-consistency readout of simulated neural activity is higher for correlated than for uncorrelated responses, thus reversing the impact of correlations on stimulus coding. Note that in the examples of this figure, model parameters did not match real data and were purely illustrative.

We then considered the readout stage of our model. Commonly, the choice is expected to follow the decoded stimulus. However, because of the apparent importance of correlations for accurate choices in our experimental data, we hypothesized that the readout of stimulus information to inform choices might utilize particular aspects of population activity imposed by correlations^1^. Intuitively, correlations imply that there is greater consistency in the neural population representations (Fig. 2e). In our model, we defined consistency as a single-trial measure of similarity between the stimuli that are decoded from features r_1_ and r_2_ separately. In our simulations, a stimulus representation in a trial was classified as consistent when features r_1_ and r_2_ both signaled the same stimulus (i.e. both features higher than average and thus both signaling s = 1, top right quadrant of panels in Fig. 2c; or both lower than average and both signaling s = −1, bottom left quadrant of panels in Fig. 2c).

We simulated binary task discrimination choices with two alternative readout models, formulated as logistic models of the dependence of choice on several features of stimulus encoding. In the first model, that we termed the consistency-independent readout, the simulated choice in each trial depended only on the stimulus decoded from the two features jointly (Fig. 2f, right panel). This case followed the traditional assumption^29^ that the choice reflects the stimulus decoded from the full population activity (Fig. 2f, readout map superimposed to left and middle panels). Note that, in our experimental data, we did not observe a perfect correspondence between the stimulus decoded from neural activity and the choice of the mouse (the fraction of times the mouse’s choice matched the decoded stimulus was 61.0% ± 0.2% in the sound localization dataset and 91.1% ± 0.1% in the evidence accumulation dataset; Supplemental Information S7). Therefore, in the model, we set the average probability that, in a given trial, the choice matched the decoded stimulus, which we termed “readout efficacy”, to a value smaller than 100%.

In the second readout model, which we termed the enhanced-by-consistency readout, the choice in each trial depended not only on the stimulus decoded from both features jointly, but also on the consistency of the stimulus decoded from the features separately (Methods) (Fig. 2i, right panel). If r_1_ and r_2_ reported consistent information about the stimulus, this readout was more likely to use the stimulus encoded in neural activity to inform the choice. This effect resulted from positive coefficients assigned to the interaction terms between the decoded stimulus and consistency (see Methods and Fig. 2i, right panel). In other words, the readout efficacy was higher when the two features were consistent (Fig. 2i, readout map superimposed to left and middle panel). Importantly, the average readout efficacy of this model was also smaller than 100%, in agreement with experimental data, and was matched to the readout efficacy of the consistency-independent readout model.

For the consistency-independent readout, correlated activity resulted in worse task performance compared to activity in which correlations were absent (Fig. 2h). This was expected because with this readout task performance is directly proportional to the level of stimulus information, with higher information resulting in higher performance. Further, noise correlations were slightly higher on simulated error trials than on correct trials (Fig. 2g), which was notably inconsistent with our experimental observations (Fig.1b, g). When considering the enhanced-by-consistency readout, noise correlations were higher in correct trials than error trials (Fig. 2j), matching our experimental findings. This readout also resulted in higher task performance in the presence of noise correlations than in the absence of correlations (Fig. 2k). Critically, this result shows that the enhanced-by-consistency readout mechanism can reconcile our experimental observations by providing a scenario in which correlations limit information, thus producing more decoding errors, while at the same time providing a benefit to task performance.

The ability of the enhanced-by-consistency readout to generate a task performance that overturns information encoding deficits was affected by how noise correlations both create consistency and limit information, which are in turn determined by the strength and structure of correlations. We explored the model parameters in depth (Extended Data Fig. 2, Supplementary Information) and found a range of parameters in which the enhanced-by-consistency readout produced higher task performance with correlations even when they limit information. This happened when correlations increased the proportion of trials with correctly decoded and consistent information more than they increased the proportion of trials with incorrectly decoded and consistent information, with respect to uncorrelated data. This condition was met when the angle between signal and noise was small but non-null, as in our experimental data (Extended Data Fig. 2b-e, Supplementary Information, Fig. 1e, j).

### Correlations and consistency in population activity contribute to behavioral choices

Together, our model results highlight the critical importance of estimating the effect of correlations not only on stimulus encoding, but also on the readout of single-trial activity to inform a choice. We therefore used our experimental measurements of neural activity to test for signatures of an enhanced-by-consistency readout. A key prediction of this readout is that the mouse’s performance should depend not only on whether stimulus encoding is correct or incorrect but also on the consistency of stimulus information across neurons and across time. In our experimental data, we defined consistency as the single-trial similarity between the stimuli that are decoded from population activity at different points in time (across-time consistency) or between the stimuli that are decoded from separate neuronal pools in the same time window (across-neuron consistency). For example, across-time consistency would be present in a trial if the population activity at time point 1 signaled the same stimulus category (e.g. s = 1) as the population activity at time point 2. We calculated the mouse’s performance for four subclasses of trials, defined by correctness and consistency of the stimulus encoded in neural activity in a given trial. In both experimental datasets, the mouse’s task performance was higher for trials with correct stimulus decoding than for incorrectly decoded trials, suggesting that the stimulus information carried by PPC neurons was used to inform behavioral choices (Fig. 3a, d, right panels). Also, the mouse’s task performance was higher for trials with consistent information across time or across neurons, supporting the idea that the consistency of neural population information was critical for accurate choices (Fig. 3a, d, left panels). Critically, in trials with correctly decoded stimulus information, the mouse’s task performance was higher when information was consistent than when it was inconsistent, both across neurons and across time. Further, in trials with incorrectly decoded stimulus information, task performance was lower on consistent trials than on inconsistent trials (Fig. 3a, d, right panels). These findings indicate that the stimulus information encoded in PPC was read out in a manner that amplified the effect of the encoded stimulus information on choice in consistent trials, both when the information was correct or incorrect.

**Figure 3:**
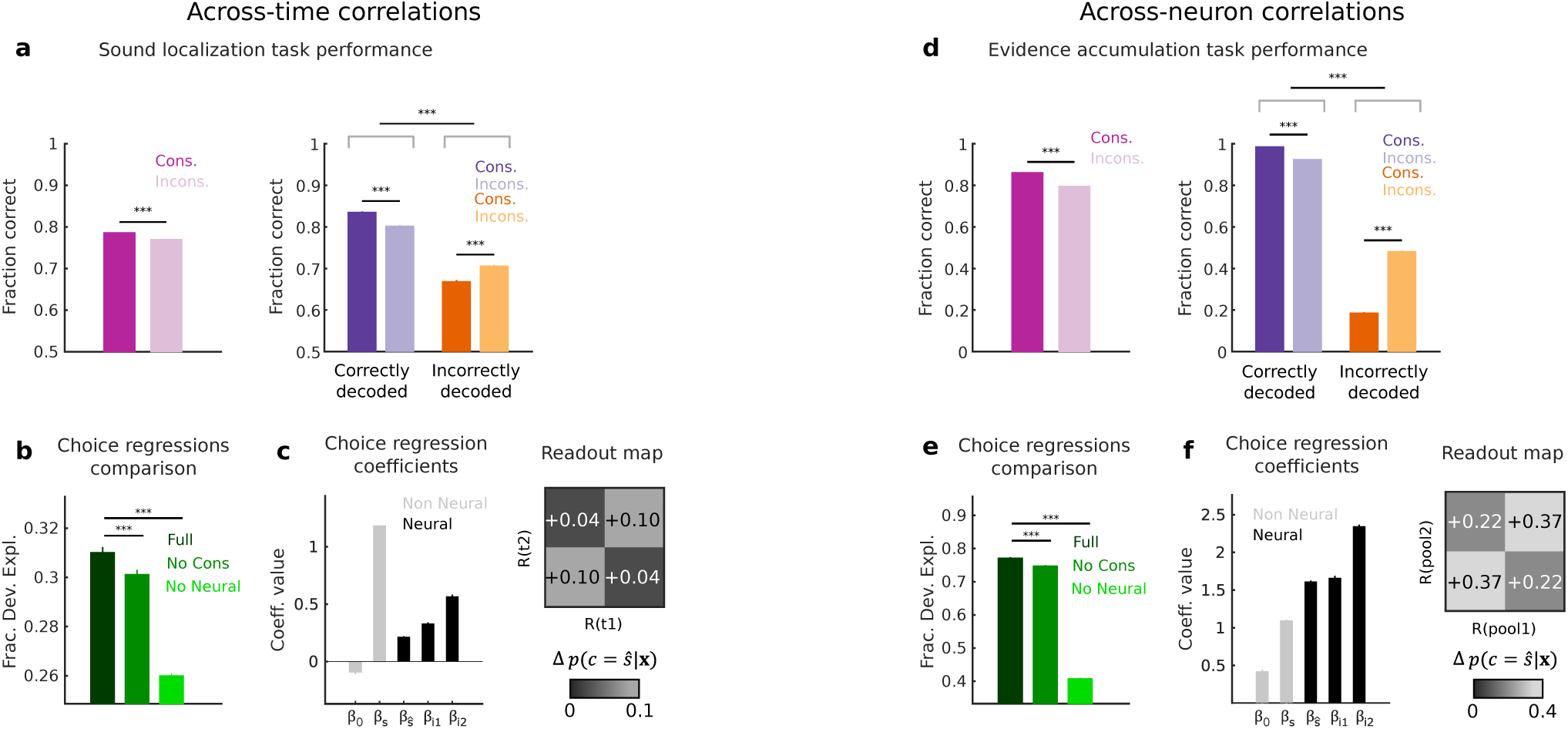
Across-time and across-neuron consistency of PPC activity influence mouse’s choices. **a**, Left: Mouse’s performance in the sound localization task is higher when neural population vectors at two different times encode the same stimulus (consistent population vectors) than when they encode different stimuli (inconsistent population vectors). Right: Mouse’s performance in the sound localization task is higher when population vectors encode the presented stimulus correctly. Furthermore, consistent population vectors are associated to average higher task performance when jointly encoding the presented stimulus correctly (left bars), and average lower task performance when jointly encoding the presented stimulus incorrectly (right bars), reflecting a consistency-dependent modulation of the animal’s readout which enhances the likelihood of encoded stimuli being converted into choices. ***P<0.001, two-sided permutation test. **b**, The readout underlying the behavioral choices of the animal is best explained by a logistic regression of choice whose predictors include, in addition to the identity of the stimulus decoded from joint PPC population vectors, its consistency across-time (Full: all predictors values extracted from recorded data; No Cons: consistency values shuffled across trials; No Neural: decoded stimulus and consistency values shuffled across trials). ***P<0.001, one-sided permutation test. **c**, Left: Best-fit coefficients of the L1-regularized full logistic regression of the mouse’s choices in the sound localization task. Right: Readout efficacy estimated from the best-fit coefficients of the full logistic regression, above and beyond the stimulus-driven baseline, for consistent and inconsistent population vectors, represented schematically as a readout map in the 2D space of population activity at two different times (see Methods). Data are mean ± s.e.m. across sessions and time point pairs with a lag difference of max 0.96 s in **a-c. d**, Left: Mouse’s performance in the evidence accumulation task is higher when neural population vectors of two randomly-selected non-overlapping neural subpopulations amongst the recorded neurons encode the same stimulus (consistent population vectors) or different stimuli (inconsistent population vectors). Right: As in **a**, for the evidence accumulation task. **e**, The readout underlying the behavioral choices of the animal is best explained by a logistic regression which predictors include, in addition to the identity of the stimulus extracted from joint PPC population vectors of non-overlapping neuronal pools, its consistency across pools. ***P<0.001, one-sided permutation test. **f**, As in **c**, for the evidence accumulation task. Data are mean ± s.e.m. across sessions, delay epochs, and 100 randomly-selected pairs of neuronal pools in **d-f**. For all panels, data are first averaged across trial splits.

To rule out that differences in a mouse’s task performance between consistent and inconsistent trials might be due to higher stimulus information in consistent trials, we sorted trials according to the stimulus information level and verified that performance in correctly (respectively incorrectly) decoded trials was still higher (respectively lower) when information was consistent across neurons or across time (Methods, Supplementary Information, and Extended Data Fig. 4c-e, g-i).

We further examined the possibility of an enhanced-by-consistency readout by developing an analytical understanding of how choices were made by the mouse. We used logistic regression to relate PPC activity to the mouse’s choices. We expressed a mouse’s choice on a given trial as a function of features of the recorded neural activity: the stimulus decoded from the full PPC population activity and its interaction with the across-time or across-neuron consistency (Methods). In addition, in the choice regression, we included a predictor for the experimenter-defined stimulus presented to the mouse and a bias term. These two terms accounted for the stimulus-related and stimulus-unrelated information carried by sources other than the recorded neurons, such as non-recorded neurons. The inclusion of these terms therefore allowed us to test how much the stimulus information in the recorded neural activity explained the mouse’s choice beyond what could be explained by other sources.

The average regression coefficients for the stimulus decoded from neural activity and the consistency-dependent interaction terms were positive (Fig. 3c, f, left), indicating that the stimulus information in neural activity, including its consistency, impacted the mouse’s choice. Positive coefficients for the consistency-dependent terms indicates that the readout of PPC activity performed similarly to the enhanced-by-consistency readout model from Figure 2: that is, the probability that the choice matched the stimulus decoded from neural activity was higher in the presence of consistency (Fig. 3c, f, right). We tested the specific contribution of the neural-based predictors in explaining the mouse’s choices by fitting the logistic regression after shuffling the values of these predictors across trials (Fig. 3b, e). Shuffling all neural-based predictors (“No Neural” choice regression in Fig. 3b, e) made it harder to predict a mouse’s choice, again demonstrating that neural activity contributed to choices. Moreover, shuffling only the neural consistency values (“No Cons” choice regression in Fig. 3b, e), while leaving intact the stimulus decoded from neural activity and the experimenter-determined stimulus, resulted in worse predictions of a mouse’s choices. Neural consistency thus seemingly contributed to the mouse’s choice.

To rule out that the modulation of the readout by consistency might just reflect differences in overall stimulus information levels between consistent and inconsistent trials, we verified that consistency provided a similar contribution to predicting a mouse’s choices when we used a more sophisticated logistic regression in which we included the magnitude of the stimulus information, instead of only the identity of the decoded stimulus (Supplementary Information and Extended Data Fig. 4b, f). Further, to control for and discount potential contributions from running-related neural activity, we verified that neural consistency also contributed to predicting choices when adding to the regression the consistency of the mouse’s running speed and direction (Supplementary Information and Extended Data Fig. 5c-e).

### The enhanced-by-consistency readout of PPC activity can overturn the information-limiting effect of correlations

Thus far, our results show that across-time and across-neuron consistency in the experimental data have an impact on a mouse’s choices. We next examined the implications of this finding for mouse task performance, in the presence or absence of experimentally-measured information-limiting correlations. Because correlations cannot be removed with current experimental approaches, we instead created a set of simulated choices using the experimentally-fit logistic choice regression from Figure 3 (Methods). We used either trials with real neural activity or trials with neural activity shuffled to disrupt across-time or across-neuron correlations as input to the experimentally-fit choice regression. As a reminder, shuffled activity had higher stimulus information (Fig. 1d, i and Fig. 4b, d, left) and lower across-time and across-neuron consistency (Fig. 4b, d, right). We used the simulated choices to provide an estimate of how well the mouse would have performed on the task with and without correlations present (Fig. 4a).

**Figure 4:**
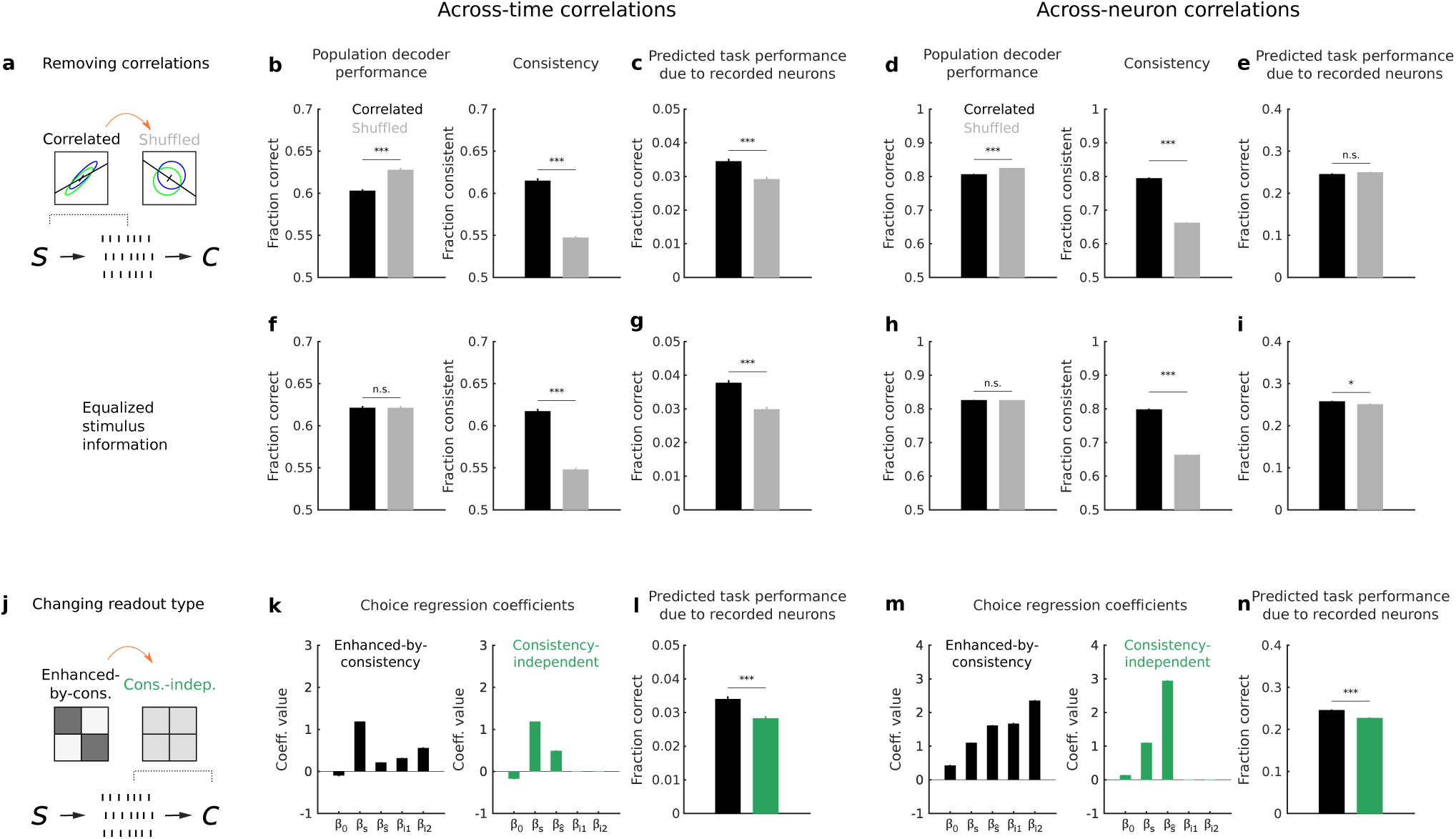
Simulated mouse’s choices show that the experimentally-fit enhanced-by-consistency readout improves task performance in the presence of information-limiting correlations in PPC. **a**, To study the impact on task performance of across-time and across-neuron noise correlations, the best-fit logistic choice regression was fed with patterns of neural activity containing the original correlations, and with artificial neural activity patterns where correlations were removed by shuffling. **b**, Left: Performance of a decoder of stimulus identity applied to joint pairs of time-lagged real recorded population activity vectors (black) and artificial shuffled population activity vectors (grey). Right: Fraction of consistent stimuli decoded separately from real and artificial pairs of time-lagged population activity vectors. **c**, Task performance attributable to the recorded neural activity, computed from choices simulated through the best-fit logistic choice regression (see Methods). Task performance was computed in the presence (black) or absence (grey) of across-time correlations between time-lagged population activity vectors. **d-e**, As in **b-c**, for across-neuron correlations. **f-g**, As in **b-c**, after subsampling trials to equalize stimulus decoder performance between real and shuffled population activity vectors. **h-i**, As in **f-g**, for across-neuron correlations. **j**, To study the impact on task performance of the consistency-dependent predictors coefficients of the best-fit choice logistic regression, the real patterns of neural activity were fed to a consistency-independent logistic regression of choice which was matched for readout efficacy with the best-fit choice regression (see Methods). **k**, Coefficients of the best-fit enhanced-by-consistency choice regression (black) and the consistency-independent choice regression (green). The latter coefficients are obtained after appropriate manipulations of the best-fit coefficients (see Methods). **l**, Task performance attributable to the recorded neural activity, computed from choices simulated through the best-fit logistic choice regression (black) or through the consistency-independent choice regression (green), in the presence of across-time correlations. **m-n**, As in **k-l**, for across-neuron correlations. Data are mean ± s.e.m. across sessions and time point pairs with a lag difference of max 0.96 s in **b-c, f-g** and **k-l**. Data are mean ± s.e.m. across sessions, delay epochs, and 100 randomly-selected pairs of neuronal pools in **d-e, h-i** and **m-n**. ***P<0.001; *P<0.05; n.s. (no statistical significance), two-sided permutation test. For all panels, data are first averaged across trial splits.

We focused on the contribution of the recorded neural population to task performance. This contribution was isolated by computing task performance from choices simulated using all predictors extracted from experimental data, and then subtracting the task performance computed from choices simulated after shuffling neural predictors values across trials (Methods). In the sound localization dataset, we estimated that the recorded ∼50 neurons increased task performance by ∼3.5% (Fig. 4c). Strikingly, the amount the recorded neurons increased task performance was greater when correlations were intact than when correlations were removed by shuffling (Fig. 4c). This is especially of note because stimulus information in the recorded neurons was lower with correlations intact (Fig. 4b, left), indicating that the enhanced-by-consistency feature of the experimentally-fit readout was able to overcome the information-limiting effect of across-time correlations and improve task performance. In the evidence accumulation data, the ∼350 recorded neurons contributed ∼25% of task performance (Fig. 4e). This estimate was similar when correlations were intact or disrupted (Fig. 4e), even though the stimulus information was lower when correlations were intact (Fig. 4d, left). This result suggests that across-neuron correlations did not hinder task performance despite having an information-limiting effect. Together, our results strongly indicate that correlations had an advantage in the readout of stimulus information that overcame or compensated for their information-limiting effects.

These results incorporate the overall impact of correlations on task performance by combining the effects of the encoding and readout stages. To analyze the specific contribution of the readout, we again simulated choices from the experimentally-fit choice regression, except we equalized the stimulus information in the correlated and shuffled populations by selecting subsets of trials having the same fraction of correctly decoded stimuli. With such matching, the correlated and shuffled trials differed only in their neural consistency (Fig. 4f, h). For both datasets, the estimated contribution to task performance of the recorded neurons was higher when correlations were intact than when they were disrupted (Fig. 4g, i). This result shows that the readout of PPC activity was more efficient in extracting information from correlated data and can thus improve task performance.

It is interesting to note that our results here indicate that the readout of stimulus information from PPC activity is suboptimal. From the ∼50 neurons recorded in the sound localization task, we were able to decode the stimulus at ∼60% correct (Fig. 4b, left). This population therefore could have increased task performance by ∼10% above chance if this stimulus information were read out optimally. However, this population only increased task performance by ∼3.5% (Fig. 4c). Similarly, for the ∼350 neurons recorded during the evidence accumulation task, we were able to decode the stimulus at ∼80% correct (Fig. 4d, left), and thus these neurons could have increased task performance by ∼30% above chance if all their stimulus information was read out. However, we estimated that these neurons only increased task performance by ∼25% (Fig. 4e). Therefore, in both datasets, the recorded neurons apparently increased task performance by a smaller amount than would have been possible if all their stimulus information was converted into choice. This result indicates that stimulus information in PPC is read out for behavior, but not optimally or entirely.

We further considered whether the experimentally-fit choice readout, which shows characteristics compatible with our enhanced-by-consistency readout model, is particularly well suited for reading out the information contained in the recorded PPC activity, or if an alternative consistency-independent readout would yield more correct choices. Our comparison was similar in spirit to that of Figure 2f-k, except using experimental data. We first generated a simulated set of choices by feeding real neural activity to the experimentally-fit regression that incorporated across-time and across-neuron consistency. We also generated a second set of simulated choices using an alternative choice regression that included only the decoded stimulus predictor, regardless of its consistency (Fig. 4j, k, m). For fairness of comparison, the coefficients for this second choice regression were selected to yield the same readout efficacy as for the experimentally-fit regression (Methods). The estimated contribution of the recorded neurons to task performance was higher with the experimentally-fit choice regression that used consistency than with the consistency-independent choice regression with matched readout efficacy (Fig. 4l, n). These experimental findings, in agreement with model predictions (Extended Data Fig. 3 and Supplementary Information), suggest that the enhanced-by-consistency readout is well suited for forming behavioral choices in the presence of information-limiting noise correlations, such as those found in PPC.

We replicated all analyses of across-neuron correlations in the sound localization dataset, that had smaller numbers of neurons per session than the evidence accumulation dataset. We found that across-neuron correlations enhanced the readout of stimulus information and task performance, even though these correlations limited information (Supplementary Information, and Extended Data Fig. 6). We could not test the effect of the across-time correlations in the evidence accumulation dataset because of challenges in dealing with time-varying stimulus evidence in that task.

Finally, in the sound localization experiments, we also had experimental data from auditory cortex. Interestingly, in auditory cortex, relative to PPC, we observed weaker noise correlations, a smaller information-limiting effect of correlations, and a lower impact of consistency on the reading out of information (Supplementary Information and Extended Data Fig. 6, 7). Therefore, sensory and association areas may differ in their levels of correlations, as we reported previously^17^, such that an enhanced-by-consistency readout may be most beneficial for reading out PPC activity and less beneficial for reading out auditory cortex activity.

## Discussion

Our results show that noise correlations limit information at the encoding stage, but surprisingly they also enhance consistency in neural codes, which improves the readout stage. The trade-off of these two effects defines the overall impact of correlations on task performance. We found that, strikingly, noise correlations can enhance task performance despite limiting the information capacity of a neural population.

Much work has emphasized that the information limiting effect of correlations in sensory areas may be a major bottleneck for behavioral performance^1,6,7^, even if the information-limiting effects of correlations that inevitably arise from common inputs may be partly reduced by anti-correlated fluctuations of excitatory and inhibitory synaptic conductances^32,33^. A largely separate set of theoretical and biophysical work has alternatively proposed that correlations improve the propagation of neural activity by mechanisms at the single neuron and network levels^10–12^. However, empirical support for a role of correlations in facilitating the readout of population information to aid behavior has been limited thus far^34,35^. First, biophysical work on signal propagation has not been connected to information coding, as it has not specified whether the transmitted activity is informative and has not distinguished between propagation of correct or incorrect information^13^, and it has been seldom connected to behavior. Second, although some studies have suggested a beneficial role of correlations by reporting higher correlation levels during correct behaviors^34,36^, these effects have not been reported when correlations limit information encoding. Further, in the analysis of our model we occasionally found conditions in which information-limiting noise correlations were stronger in correct trials, without them having a positive effect on behavioral performance, suggesting that reports of higher correlations in correct trials is indicative that consistency affects readout, but it is not sufficient to imply a behavioral performance advantage of correlations (see Extended Data Figure 2 and Supplemental Information S8). Thus, from previous work it may be challenging to conclude whether the advantages of correlations for signal propagation can overturn their information-limiting effect.

Here we developed a formalism that bridges between the two separate research streams of information encoding and signal propagation, to address how the negative encoding effects and the positive readout effects of correlations intersect. Remarkably, in our data the advantages of correlations for signal propagation were large enough to compensate and even overcome the negative encoding effects. We unraveled a readout computation that could not be inferred from previous biophysical activity propagation studies: that when correlations are information limiting the interaction between the readout and encoding of stimulus information is such that it amplifies correctly decoded sensory information more than incorrectly decoded information.

Our approach provides a novel and generally applicable framework to parametrically dissect the contribution of correlated neural activity to perceptual behaviors. We anticipate that this approach can be applied to different tasks and brain areas. Our initial observations comparing primary sensory and association cortices suggests that the best trade-off between the effect of correlations on encoding and readout may vary depending on the area. In sensory cortices, low correlations and a readout insensitive to consistency may be advantageous to encode rapidly changing and high dimensional sensory features, in order to represent many features of the sensory environment regardless of if they are used for the immediate behavior at hand. This view is compatible with reciprocal relationships between noise correlation levels and behavioral performance in sensory cortices^1^. In contrast, association cortices that are closer to behavioral output may only need to encode a moderate amount of behaviorally relevant sensory information but this information should have a strong impact on behavior. In these areas, the price to be paid for reducing encoded information is smaller and the price to be paid for losing signal in the propagation to behavior is higher. Thus, in association areas, the best tradeoff may be to have some redundancy in the neural representation coupled with a readout mechanism that uses this redundancy to enhance signal propagation to inform choice, as we found here. Further, we anticipate that our general formalism will allow us to design direct causal tests of the actual readout used in the brain during perceptual discrimination tasks. We propose that our formalism together with holographic perturbations^37^ will allow us to test the behavioral role of correlations and consistency across neurons and across time.

Noise correlations can reflect interactions between cells, shared covariations due to common inputs, general fluctuations in behavioral state or network excitability, or even variations with stimuli within the same category^2^. This shared variability, independently of its origin, acts as noise (that cannot be reduced by integrating information over more cells or longer times) from the decoding point of view, but helps signal propagation by generating more consistent neural representations. Thus, our conclusions hold independently of the specific biophysical origins of the observed noise correlations.

Most studies of neural coding implicitly or explicitly assume that the readout of sensory information is optimal and interpret neural codes with higher sensory information as being more relevant for perception^6,28,29^. Part of the reason for this assumption is that the presence and shape of non-optimality are unknown. If the readout is not optimal, then neural codes with higher information are not necessarily the most relevant ones for perception. Our study suggests (see Supplemental Information) that a sub-optimality in the readout of our PPC data exists, because stimulus information in neural activity seems not to be used in its entirety to produce accurate behavioral choices, as also predicted by other studies^38,39^. Our work provides a measure of both the nature of readout non-optimality and its implication for the behavioral relevance of a neural code. Previous work on the optimality of readouts has examined whether a decoder of correlated population activity could be trained sub-optimally to decode separately single-cell data and then join together their evidence^40–42^. Several of these studies have shown that even relatively simple decoders trained on single cells can decode stimulus information from population activity, so that correlations among cells do not greatly complicate the extraction of information. However, unlike ours, these previous studies did not examine the effect of correlations on determining behavioral choices. Taken together, emerging evidence suggests that correlations do not necessarily greatly complicate the decoding of sensory information and may offer advantages for turning sensory information into appropriate behavioral choices.

## Acknowledgements

We thank E. Piasini for his early contribution to our analyses, members of our laboratories for helpful discussions, G. Iurilli, C. Kayser, E. Piasini and J. Drugowitsch for feedback on the manuscript. This work was supported by NIH grants from the NIMH BRAINS program (R01 MH107620), NINDS (R01 NS089521), the BRAIN Initiative (R01 NS108410 and U19 NS107464), and the Fondation Bertarelli.

## Author Contributions

SP and CDH conceived and supervised the study. CAR and ASM acquired the experimental data. MV and GP performed the computational work. SP, CDH, MV and GP wrote the manuscript, with feedback from all authors.

---

**Extended Data Figure 1:**
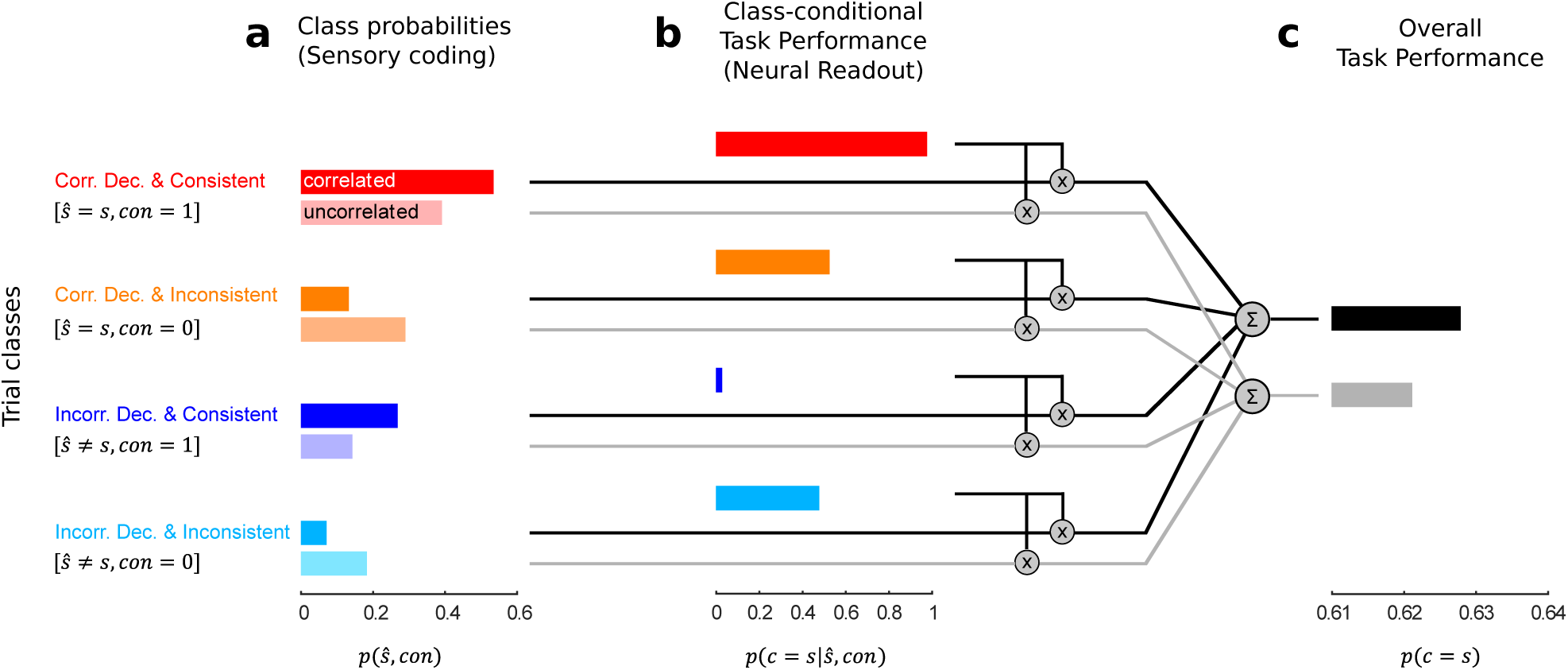
Task performance as a sum of encoding-by-readout terms depending on correctness and consistency of encoded stimulus information. **a.** Probabilities for a set of n=200 trials drawn separately from the correlated (dark colors) and uncorrelated (light colors) distributions sketched in Fig. 2c to belong to one of the four trial classes defined by correctness of decoded stimulus and consistency. Correlations overall decrease the fraction of correctly decoded trials (red + orange bars). At the same time, correlations increase the fraction of consistent trials which encode the correct stimulus identity (red), but also the fraction of consistent trials which encode the incorrect stimulus identity (blue). **b**, Average task performance for each of the four classes of trials in **a**, determined by the enhanced-by-consistency readout model coefficients sketched in Fig. 2i. The enhanced-by-consistency readout is characterized by close-to-random task performance for inconsistent trials, close-to-totally-correct choices for correctly decoded consistent trials and close-to-totally-wrong choices for incorrectly decoded consistent trials. **c**, Overall task performance depends on both on the stimulus decoding correctness and consistency of trials characterized by a given correlation structure and on the class-conditional task performance imposed by the readout model. Note that under hypothesis of enhanced-by-consistency readout it is possible for correlations that decrease the overall fraction of correctly decoded trials to increase the overall fraction of correct choice trials (with respect to the uncorrelated equivalent).

**Extended Data Figure 2:**
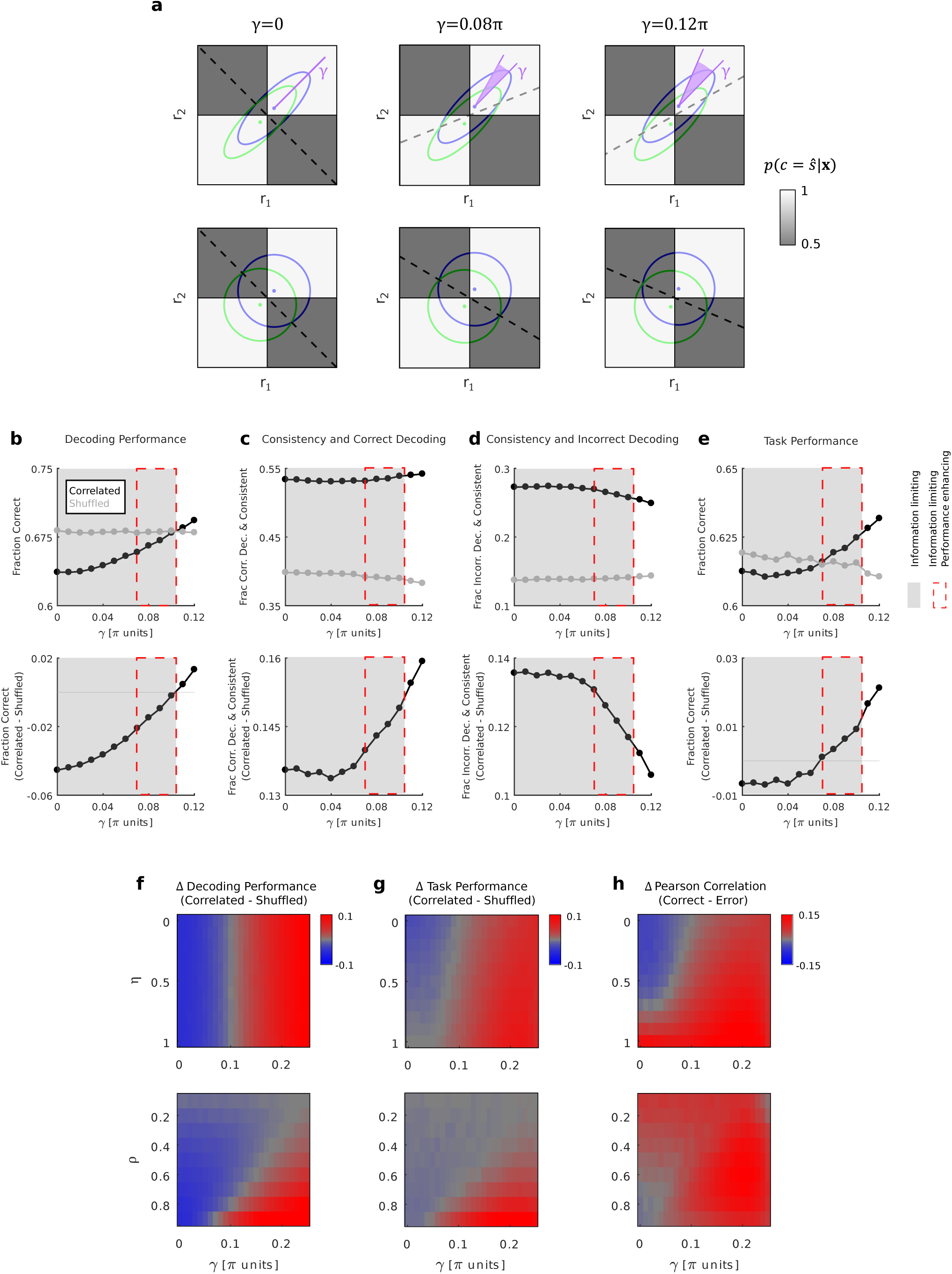
Exploration of the parameter space of the two-feature encoding readout model, to investigate the conditions in which an enhanced-by-consistency readout produces an advantage for task performance for data characterized by information-limiting correlations. **a**, Example Gaussian response distributions along two neural features (r_1_,r_2_) to two stimuli (s=1: blue, s=-1: green) at different degrees of alignment between signal and noise (γ) for correlated features (top, *ρ* = 0.8), and the corresponding shuffled distributions (bottom, *ρ* = 0). Ellipses indicate 95% confidence intervals. The dashed black line indicates the optimal decoding boundary. Note how, when increasing γ, the correlated response distributions to the two stimuli become less overlapped. **b**, Top: Performance of a linear decoder of stimulus identity applied to correlated (black) and uncorrelated (grey) responses for a range of γ values. The shaded grey region indicates values of γ for which correlations are information limiting. **c**, Top: The fraction of trials being correctly decoded and carrying consistent stimulus information increases at increasing γ for correlated but not for uncorrelated responses. **d**, Top: The fraction of trials being incorrectly decoded and carrying consistent stimulus information decreases at increasing γ for correlated but not for uncorrelated responses. **e**, Top: Task performance predicted by applying the enhanced-by-consistency readout model in Fig. 2i to correlated and uncorrelated responses for a range of γ values. The dashed red rectangle indicates values of γ for which information limiting correlations are advantageous for task performance. **b-e**, Bottom: Difference between correlated and uncorrelated responses, for the quantities shown in the top panels. **f**, Difference between stimulus decoding performance from correlated and uncorrelated responses for different combinations of model parameters values (γ: alignment between signal and noise. *η*: strength of consistency modulation in the readout. *ρ*: noise correlations strength). **g**, Difference between task performance predicted by applying the enhanced-by-consistency readout model in Fig. 2i to correlated and uncorrelated responses for different combinations of model parameters values. Note that there exist a range of model parameters for which correlations at the same time decrease decoding performance and improve task performance. **h**, Difference in Pearson correlation between correct choice and incorrect choice trials from correlated responses for different combinations of model parameters values. Where not otherwise specified, *ρ* = 0.8 and *η* = 0.7 in **b-h**.

**Extended Data Figure 3:**
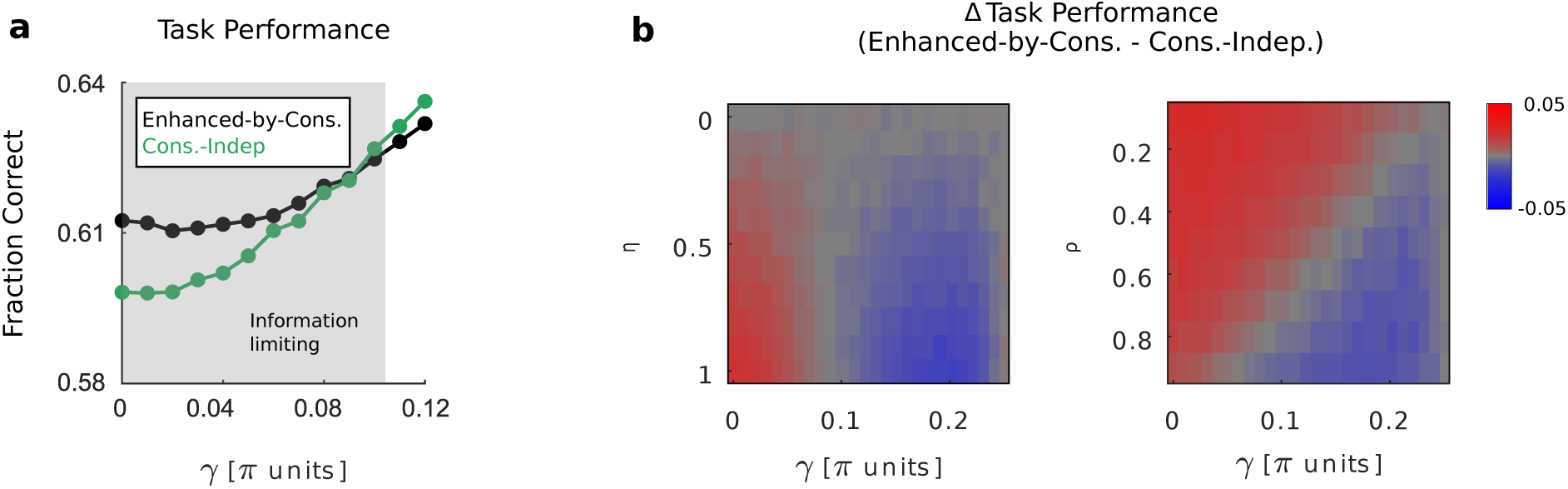
Exploration of the parameter space of the two-feature encoding readout model, to investigate the conditions in which an enhanced-by-consistency readout produces an advantage for task performance over a consistency-independent readout with matched readout efficacy. **a**, Task performance predicted by applying the enhanced-by-consistency readout in Fig. 2i (black) or a consistency-independent readout with matched readout efficacy (green, see Methods) to simulated correlated responses (as in Extended Data Fig. 2a, top), for different degrees of alignment between signal and noise (γ). The shaded grey region indicates values of γ for which correlations are information limiting. **b**, Difference between task performance predicted by applying the enhanced-by-consistency readout and the one predicted by applying the consistency-independent readout to correlated responses for different combinations of model parameters values (γ: alignment between signal and noise. *η*: strength of consistency modulation in the readout. *ρ*: noise correlations strength). Where not otherwise specified, *ρ* = 0.8 and *η* = 0.7 in **a-b**.

**Extended Data Figure 4:**
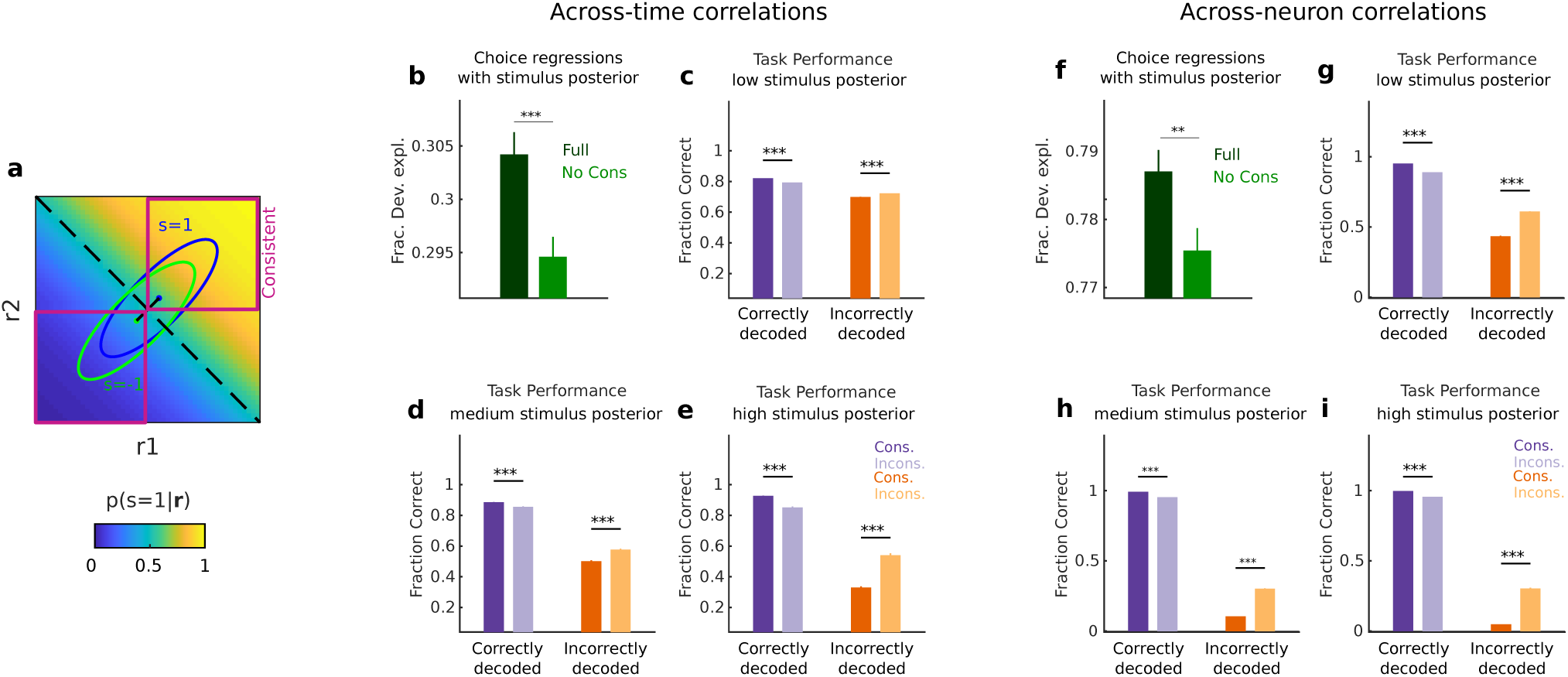
The effect of neural consistency on the mouse’s single trial choices cannot be explained by higher stimulus information associated to consistent neural representations. **a**, Schematic example showing response distributions along two neural features (r_1_, r_2_) to two stimuli (*s* = −1: green, *s* = 1: blue), which are best separated by the black dashed line, which represents the decoding boundary of a linear decoder trained on the observed responses. The background color encodes the linear decoder posterior probability that stimulus s=1 has occurred given the observation of the neural response **r** = (r_1_, r_2_). Intuitively, the farther neural response ***r*** is from the decoding boundary, the farther *p*(*s* = 1|***r***) is from 0.5, the more “informative” ***r*** is about the stimulus. Note that, in the depicted example, consistent trials have on average higher posterior probability than inconsistent trials, which might represent a confounder for the effect of consistency on mouse’s choices. **b**, The readout underlying the choices of the animal is best explained by a choice regression that includes, in addition to the posterior probability of the stimulus extracted from joint PPC population vectors, its consistency across-time (Full: all predictors values, including stimulus posterior probability and posterior probability consistency, extracted from recorded data; No Cons: posterior probability consistency values shuffled across trials). ***P<0.001, one-sided permutation test. **c-e**, In order to control for possible confounding effects of the posterior probability in determining the behavioral relevance of across-time consistency, the computation in Fig 3b-c was repeated considering only subsets of trials having low (|*p*(*s* = 1|***r***) − 0.5| < 0.16), medium (|*p*(*s* = 1|***r***) − 0.5| > 0.16 ⋀ |*p*(*s* = 1|***r***) − 0.5| < 0.32) or high (|*p*(*s* = 1|***r***) − 0.5| > 0.32) posterior. Consistent population vectors result in average higher task performance when jointly encoding the presented stimulus correctly (left bars), and average lower task performance when jointly encoding the presented stimulus incorrectly (right bars), even when subsets of trials having approximately the same posterior are used. ***P<0.001, two-sided permutation test. **f-i**, as **b-e**, for across-neuron correlations. Data are plotted as mean ± s.e.m. across sessions and time point pairs with a lag difference of max 0.96 s in **b-e**. Data are plotted as mean ± s.e.m. across sessions, delay epochs, and 100 randomly-selected pairs of neuronal pools in **f-i**. For panels **b-i**, data are first averaged across trial splits.

**Extended Data Figure 5:**
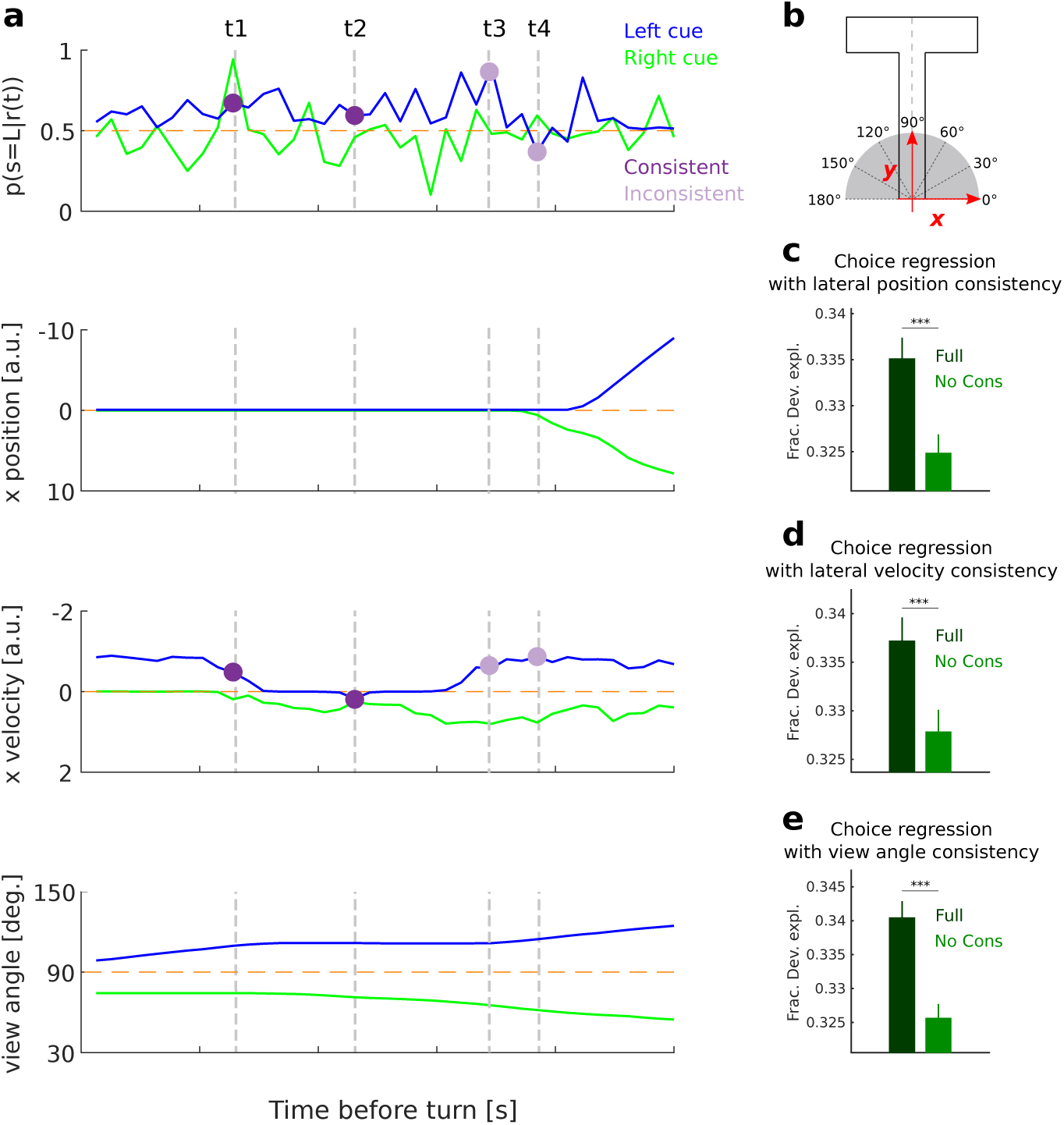
The role of neural consistency in the readout of PPC activity is not due to the consistency of measured behavioral parameters. **a**, The across-time evolution of the decoder posterior probability of left stimulus presentation given the recorded PPC population activity (sound localization task dataset) is shown along with the corresponding across-time evolution of a selection of three concurrently-measured behavioral parameters (lateral position, lateral velocity, view angle), for an example left (blue) and right (green) cue trial. Example time point pairs with consistent (t_1_-t_2_, dark purple) or inconsistent (t_3_-t_4_, light purple) neural information are highlighted. Neural consistency is not necessarily associated to behavioral consistency (e.g. when considering lateral running speed, t_1_-t_2_ are behaviorally inconsistent while t_3_-t_4_ are behaviorally consistent). **b**, Schematic representation of the virtual T-maze with corresponding x-y coordinates labelling and mouse’s view angle (for a mouse oriented along the y axis). **c-e**, The readout underlying the mouse’s choices is best explained by a choice regression that includes, in addition to the consistency of measured behavioral parameters (**c**: lateral position, **d**: lateral velocity, **e**: view angle), the consistency of the stimulus identity extracted from neural activity (Full: all predictors values, including neural and behavioral consistency, extracted from recorded data; No Cons: neural consistency values shuffled across trials). For panels **c-e**: data are plotted as mean ± s.e.m. across sessions and time point pairs with a lag difference of max s; data are first averaged across trial splits; ***P<0.001, one-sided permutation test.

**Extended Data Figure 6:**
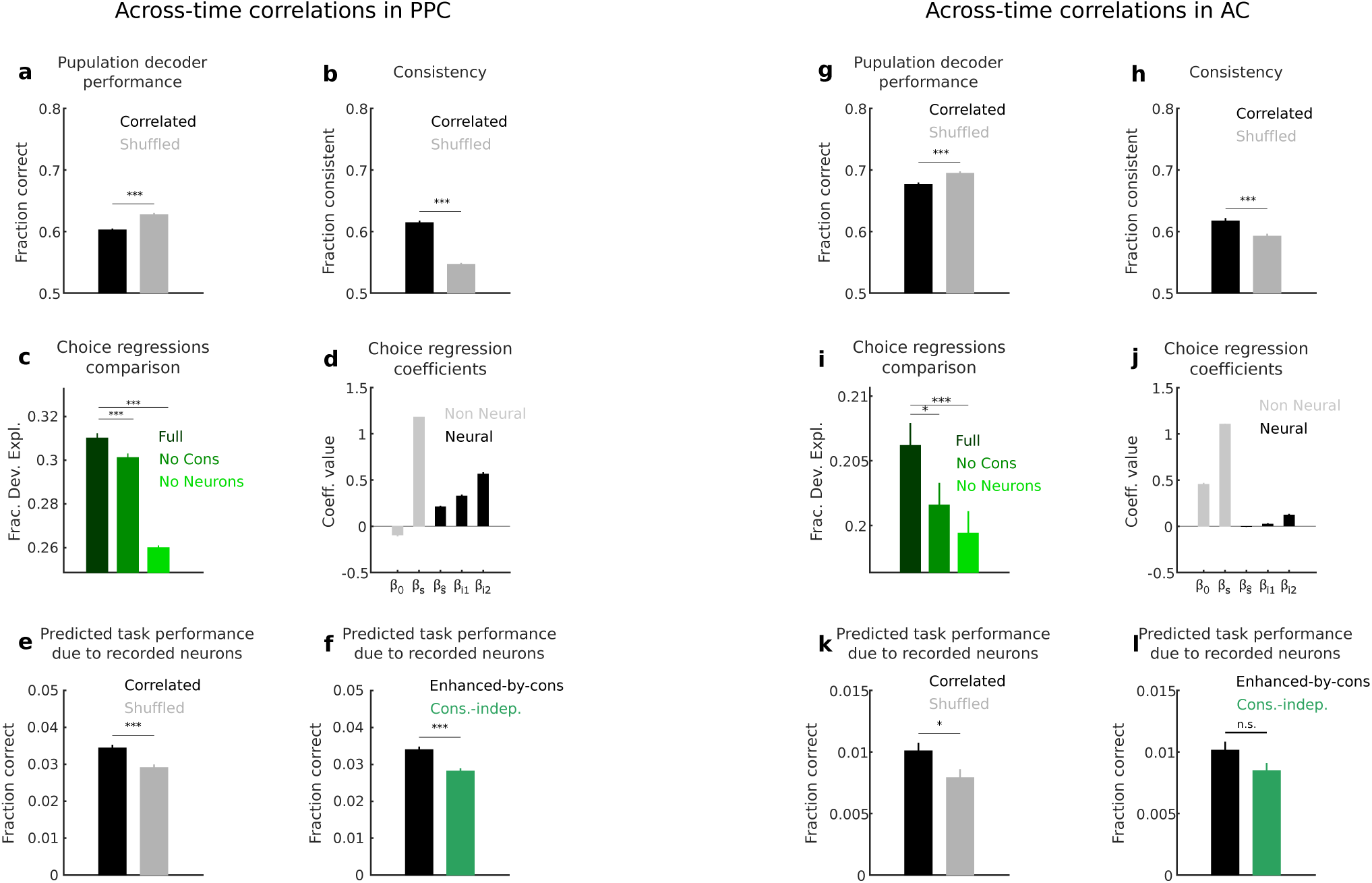
Across-time consistency of stimulus information in AC does not benefit task performance as it does in PPC. **a-f**, Summary of the main results of the analysis of across-time correlations in PPC activity (sound localization task dataset), reported in Fig. 3, 4, useful for comparison with the corresponding results of analysis of across-time correlations in AC activity (sound localization task dataset), reported in **g-h. g**, Performance of a decoder of stimulus identity applied to joint population activity at two different time points is higher in AC than in PPC. Across-time correlations limit the encoding of stimulus information in AC just like in PPC. **h**, Across-time correlations in AC increase the consistency of encoded stimulus information less than across-time correlations in PPC. ***P<0.001, two-sided permutation test. **i**, Across-time consistency of stimulus information encoded in AC does not improve substantially the performance of the logistic regression in predicting the animal’s choices. ***P<0.001; *P<0.05, one-sided permutation test. **j**, Best-fit beta coefficients of the full choice regression, showing that AC neural predictors are assigned very low weights in the readout. **k**, Task performance that can be attributed to the recorded neurons is much lower in AC than in PPC (∼1% in AC, ∼3.5% in PPC). Correlations in AC activity happen to enhance task performance, but the effect is very small. **l**, Task performance which is attributable to the recorded AC neural activity would not be substantially different if the behavioral readout was consistency-independent. *P<0.05; n.s. (no statistical significance), two-sided permutation test. Data are mean ± s.e.m. across sessions and time point pairs with a lag difference of max 0.96 s. Data are first averaged across trial splits.

**Extended Data Figure 7:**
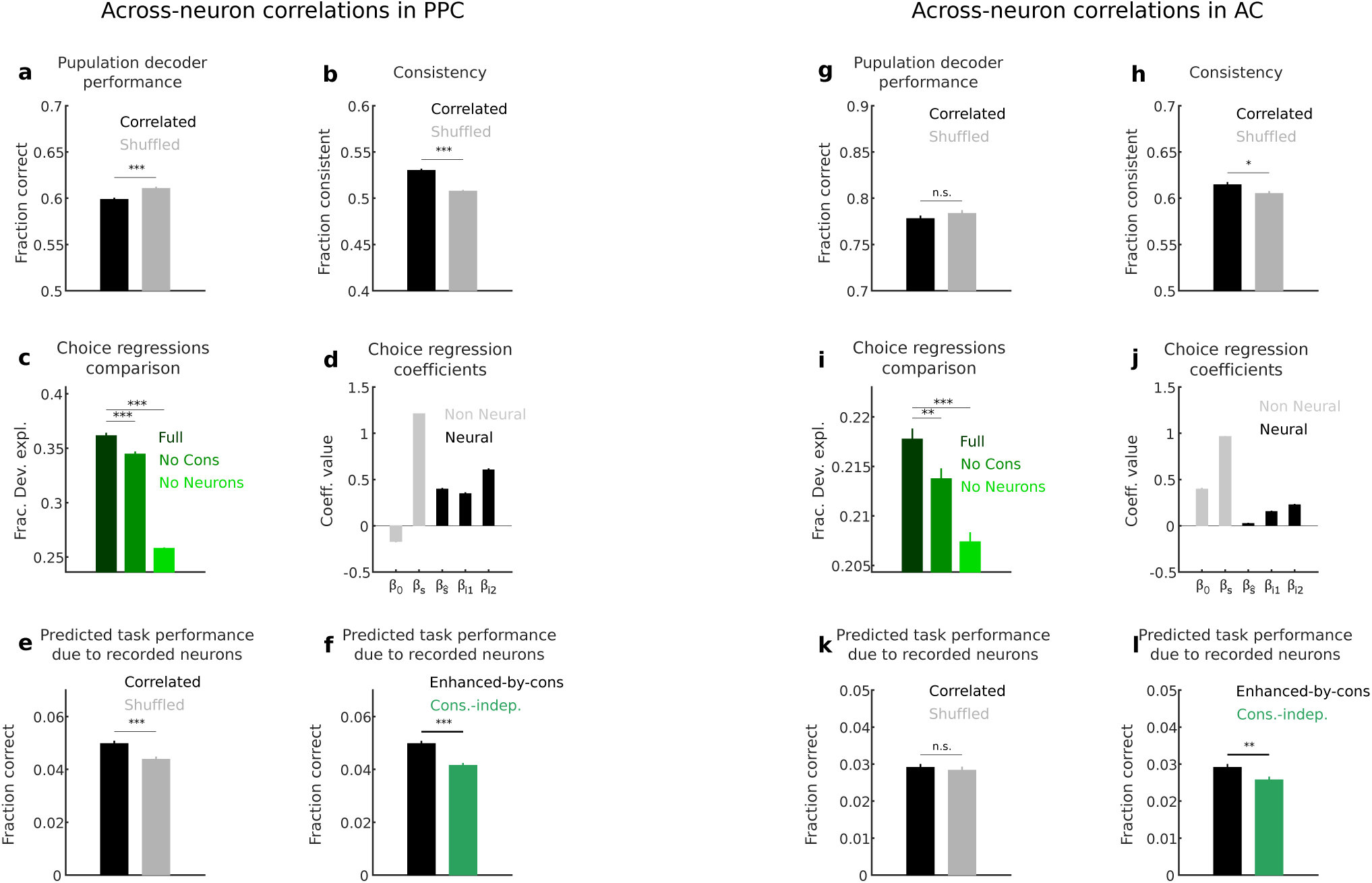
In the sound localization dataset, across-neuron correlations in PPC are information limiting and provide an advantage for task performance. **a**, Across-neuron correlations in PPC limit the encoding of stimulus information during the sound localization task. **b**, Across-neuron correlations increase the consistency of encoded stimulus identity across neuronal subpopulations; however, the effect is less strong than that of across-time correlations (compare with Extended Data Fig. 6b). ***P<0.001, two-sided permutation test. **c**, The readout underlying the behavioral choices of the animal in the sound localization task is best explained by a choice regression that includes, in addition to the identity of the stimulus decoded from joint PPC population vectors, its consistency across neurons. ***P<0.001, one-sided permutation test. **d**, Best-fit beta coefficients of the full regression (which best explains the behavioral choices of the animal), showing that the across-neuron consistency plays a relevant role in the readout. **e**, Task performance in the sound localization task that can be attributed to the recorded neurons benefits from the presence of correlations across neurons in PPC. **f**, Task performance in the sound localization task that can be attributed to the recorded neurons would be lower if the readout of PPC activity was independent from consistency across-neurons. ***P<0.001, two-sided permutation test. **g-l**, Similarly to the case of across-time correlations (see Extended Data Fig. 6g-l), across-neuron correlations in AC are weak and so is their effect on stimulus information encoding (**g, h**) and predicted task performance (**k**). Across-neuron consistency seem to play a more important role in readout than across-time consistency in AC (**i, j, l**). Data are mean ± s.e.m. across sessions, time windows with a max duration of 0.96 s and 100 randomly-selected pairs of neuronal pools. Data are first averaged across trial splits.

## Methods

No statistical methods were used to predetermine sample size. The experiments were not randomized. The investigators were not blinded to allocation during experiments and outcome assessment.

### Note on present study

This study represents an independent analysis of experiments described previously^17,19^. Although the raw data are the same, the questions investigated and analytical tools are different, focusing here on the analysis of the role of across-time and across-neuron correlations on information coding and perceptual discrimination performance.

### Subjects, behavioral task and two-photon imaging

A brief summary of the experimental procedures is provided here. A more detailed description can be found in the original works^17,19^. All experimental procedures were approved by the Harvard Medical School Institutional Animal Care and Use Committee.

Both experiments made use of a modified version of a visual virtual reality system that has been described previously^44^. Head-restrained mice ran on a spherical treadmill, while images of a virtual maze were projected on a half-cylindrical screen. Forward/backward translation in the maze was controlled by treadmill changes in pitch, and rotation in the virtual environment was controlled by roll of the treadmill. The virtual maze was constructed using the Virtual Reality Mouse Engine (VirMEn) in MATLAB^45^.

#### Sound localization task dataset

Imaging data were acquired from five male C57BL/6J mice (The Jackson Laboratory), aged 6-8 weeks at the initiation of behavioral task training. Imaging began 4-6 weeks after viral injection and was continued for 4-12 weeks.

In the final version of the task that was used during imaging, mice ran down the stem of the virtual T-maze, while sound stimuli were delivered from 8 possible locations (−90°, −60°, −30°, −15°, +15°, +30°, +60°, +90°) using four electrostatic speakers positioned in a semicircular array, centered on the mouse’s head. The sound stimulus was activated when the mouse passed an invisible spatial threshold at ∼10 cm into the T-stem and originated from one of eight possible locations. The stimulus was repeated after a 100 ms gap; repeats continued until the mouse reached the T-stem. Task difficulty was modulated by the direction of the incoming stimulus, with the −90°/+90° trials being the easiest ones and the −15°/+15° trials being the most difficult ones. In order to receive a reward (4 µl water), mice had to judge the location of sound stimuli to be either on the left or right, and to report their decisions by turning left or right at the T-intersection. A “reward tone” was played as the water reward was delivered on correct trials (when the mouse had reached ∼10 cm into the correct arm of the T-maze), and a “no reward tone” was played when the mouse reached ∼10 cm into the incorrect arm on error trials. The inter-trial interval was 3 s on correct trials and 5 s on error trials. Mice performed ∼200 trials (range, 125–251) in a typical session over approximately 45–60 minutes.

Imaging was performed on alternating days from AC and PPC on the left hemisphere (PPC centered at 2 mm posterior and 1.75 mm lateral to bregma; AC centered at 3.0 mm posterior and 4.3 mm lateral to bregma). In a given session, ∼50 neurons (range, 37-69) were simultaneously imaged using a two-photon microscope (Sutter MOM) operating at 15.6 Hz frame rate and at 256 × 64 pixel resolution (∼ 250 µm × 100 µm). Imaging data were acquired at depths between 150 and 300 µm, corresponding to layers 2/3. Seven AC fields of view and seven PPC fields of view from five mice were used for analysis.

#### Evidence accumulation task dataset

Imaging data were acquired from five male C57BL/6J mice (The Jackson Laboratory), aged 8–10 weeks at the at the initiation of behavioral task training. Imaging began at least 4 weeks after viral injection and was continued for up to 12 weeks.

In the final version of the task that was used during imaging, mice run down the stem of a virtual T-maze with predominantly gray walls, encountering six visual cues (white wall segments with black dots) at fixed locations. Each cue could appear on either the left or right wall, and only one cue was visible at a time. To receive a reward (4 µl 10% sweetened condensed milk), mice had to determine whether more cues were presented on the left or the right and, after a short stretch of maze without additional cues (90 cm), turn at the T-intersection toward the direction that had more cues (left for 6-0, 5-1, 4-2 trials; right for 2-4, 1-5, 0-6 trials). Task difficulty was modulated by varying the difference between the number of left and right cues (net evidence). The sequence of cues was determined randomly for each trial of a given net evidence. On trials having zero net evidence (3-3 trials), the rewarded location was selected randomly. Following the completion of the trial, the screen changed to black for the duration of the inter-trial interval (2 s for correct choice and 4 s for incorrect choice). Mice performed ∼300 trials (range, 231–414) in a typical session over approximately 45–60 minutes.

Imaging data were acquired from the left PPC (PPC centered at 2 mm posterior and 1.75 mm lateral to bregma). In a given session, ∼350 neurons (range, 188-648) were simultaneously imaged using a custom-built two-photon microscope operating at ∼30 Hz frame rate and at 512 × 512 pixel resolution (∼700 µm × 700 µm). Imaging data were acquired at depths between 100 and 200 µm below the dura. Eleven fields of view from five mice were used for analysis.

### Imaging data processing

Imaging frames were concatenated for motion correction, cell body identification, extraction of fluorescence traces and estimation of relative spike rates. In brief, after motion correction^46^, correlations in fluorescence time series between pixels within ∼60 µm were calculated. Fluorescence sources (putative cells) were identified by applying a continuous-valued eigenvector-based approximation of the normalized cuts objective^47^ to the correlation matrix, followed by k-means clustering segmentation, yielding binary masks for all identifiable fluorescence sources. To estimate potential neuropil contamination, the cell body fluorescence signal was regressed against the signal from surrounding pixels during the imaging frames when the cell of interest was not active, and then neuropil contamination was removed during the ΔF/F calculation by subtracting a scaled version of the neuropil signal from the cell body signal. All fluorescence traces were deconvolved to estimate the relative spike rate in each imaging frame, which was expressed in arbitrary units given that the estimate is relative to baseline activity without perfect knowledge of the fluorescence change associated with a single spike^48^.

### Data inclusion and task epoch selection for encoding and readout analyses

#### Sound localization task dataset

Population activity data from the sound localization task dataset were used for the analysis of across-time correlations in PPC reported in the main text and for additional analyses of across-time correlations in AC and across-neuron correlations in PPC and AC reported in Supplementary Information.

For the analysis of across-time correlations in PPC, population activity data were temporally aligned to the imaging time frame of the turn, defined as the frame in which the mouse entered the short arm of the maze. Since it is reasonable to assume that the animal computes its choice after the stimulus presentation but before the turn, the analysis focused on the 39 frames preceding the turn frame (this number of frames was chosen because it covered the maximum portion of the pre-turn period that was commonly available across all recording sessions). One of the seven PPC recording sessions used in our previous published work^17^ was excluded due to the large unbalance of left/right stimuli that were presented to the mouse across trials in that session, which would result in too few trials available for following analyses (see “Analysis of stimulus encoding and consistency”).

For the analysis of across-time correlations in AC, population activity data were temporally aligned to the imaging time frame of the first auditory stimulus presentation, and the analysis focused on the 50 frames after that frame (this number of frames was chosen because it covered the maximum portion of the post-stimulus period that was commonly available across all recording sessions). AC neural data aligned to the turn did not encode a sufficient amount of stimulus information for following analyses. One of the seven AC sessions used in our previous published work^17^ was excluded due to the large unbalance of left/right stimuli that were presented to the mouse across trials in that session.

For the analysis of across-neuron correlations in PPC, population activity data were temporally aligned to the imaging time frame of the turn and averaged over ten turn-aligned time windows of increasing length, comprising the {4:8: … :36:39} frames preceding the turn imaging frame. The same six sessions used for the analysis of across-time correlations in PPC were used.

#### Evidence accumulation task dataset

Population activity data from the evidence accumulation task dataset were used for the analysis of across-neuron correlations in PPC reported in the main text. PPC population activity data were first grouped into spatial bins (3.75 cm/bin) covering the whole length of the T-maze (long and short arm) by averaging population activity in each bin (2 or 3 imaging frames per bin per trial). Population activity data were then further averaged over epochs of 4 spatial bins each (about 200 ms). We considered the same 10 epochs that were defined in the original work^19^. We plotted results corresponding to the population activity data recorded in the Early Delay and Late Delay epochs. During the delay epochs, the cue presentation was completed but the animal had not yet committed to a final turn (more precisely, these epochs correspond to the four spatial bins beginning respectively 15 and 37.5 cm after the offset of the final cue). Therefore, it is reasonable to assume that these epochs are involved in the formation of the animal’s decision. All 11 sessions in the original work^19^ were used. We discarded trials with zero net evidence (<10% trials in 2/11 sessions).

### Selectivity of single cells to stimulus category

For Fig. 1c, h, we computed the selectivity of single cells to stimulus category. Stimulus category was defined as a binary variable (left/right). For the sound localization task, stimulus category corresponds to the direction of incoming auditory stimuli. Each stimulus category comprises four different sound locations (left: −90°, −60°, −30°, −15°; right: +15°, +30°, +60°, +90°). For the evidence accumulation task, stimulus category corresponds to the side of the maze where the majority of the visual cues were presented (left: 6-0, 5-1, 4-2 trials; right: 2-4, 1-5, 0-6 trials).

The selectivity index^15^ (SI) was quantified as:

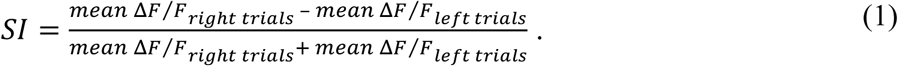

Cells that had a selectivity index greater than zero were considered right-preferring cells and those that had a selectivity index less than zero were considered left-preferring cells.

### Pairwise noise correlations

We quantified across-time and across-neuron pairwise noise correlations separately for trials in which mice made correct or error choices. In order to discount possible differences in correlation related to trials numerosity, we subsampled trials to equalize the number of correct and error trials in each recorded session. The subsampling procedure was repeated 20 times, and results were averaged over the 20 subsamples.

#### Across-time correlations

We quantified across-time pairwise correlations by computing, for each pair of recorded neurons and for each pair of time points in the trial epoch selected for analyses (see “Data inclusion and task epoch selection for encoding and readout analyses”), the Pearson correlation between the activity of neuron 1 at time t_1_ and the activity of neuron 2 at time t_2_ across trials sharing the same stimulus category. Results were averaged first across all time point pairs sharing the same lag between each other, then across stimuli and finally across trials subsamples.

#### Across-neuron correlations

We quantified across-neuron pairwise correlations by computing the Pearson correlation for each pair of neurons recorded in a single session, across trials sharing the same stimulus category. Results were averaged first across stimuli and then across trials subsamples.

### Analysis of stimulus encoding and consistency

For encoding and consistency analyses, we considered information about stimulus category (see “Selectivity of single cells to stimulus category”).

Information about stimulus category carried by population activity was extracted by decoding the most likely stimulus category presented to the animal in each trial using a C-support vector classifier (C-SVC) with a linear basis function kernel^49,50^. The C-SVC was implemented with custom MATLAB software, by building on the libsvm library^51^. For each imaging session, we first subsampled trials randomly such that the left/right stimulus categories were equally represented in the data (sound localization task dataset: no more than 13% of removed trials per session; evidence accumulation task dataset: no more than 15% of removed trials per session). Then, we randomly split the remaining trials 10 times into 50/50 training/testing sets, such that left and right stimulus categories were equally represented in both training and testing sets. For each trial split, we trained the C-SVC on the training set and we tested on the test set, which was left out of the fitting procedure. The regularization hyperparameter (C) was selected by maximizing the 3-fold cross-validated decoding accuracy in the training set. For the analyses that required computing a posterior probability of the decoded stimulus given the observed population activity, we used Platt scaling to calibrate posterior probabilities on the binary outputs of the C-SVC^52^.

#### Across-time consistency

For the across-time correlations analysis, we decoded the most likely stimulus from the population activity at each time point in the trial epoch selected for analyses (see “Data inclusion and task epoch selection for encoding and readout analyses”) and from the concatenated population activity of every pair of time points. Any two time points were defined to be consistent if the stimuli decoded from the two time points individually coincided.

#### Across-neuron consistency

For the across-neuron correlations analysis, we first split the neuronal population recorded in each session into two randomly-selected, equally-sized, disjoint pools of neurons. We then decoded the most likely stimulus from the population activity of each pool individually and from the concatenated population activity of both pools (note that this corresponds to the activity of the whole recorded neuronal population). Two pools were defined to be consistent if the stimuli decoded from the two pools individually coincided. The random assignment of the neurons to the two pools was repeated 100 times.

### Quantifying the angle between the signal and noise axes

We quantified, in the neural population response space, the angle *γ* between the direction of maximum stimulus variation (signal correlations axis) and the direction of maximum noise variation (noise correlations axis)^24,25^. The signal correlations axis was computed as the slope of the linear regression of the trial-averaged responses at fixed stimulus. The noise correlations axis was computed as the slope of the linear regression of the trial-by-trial neural responses at fixed stimulus, averaged across stimuli. *γ* takes values between 0 and *π*/2.

#### Across-time correlations

For across-time correlations, multidimensional population responses at a given time point were first reduced to scalars by computing the projection along the direction of maximum stimulus variation in the space spanned by the population responses at the selected time point. Then, for each pair of time points in the trial epoch selected for analyses (see “Data inclusion and task epoch selection for encoding and readout analyses”), *γ* was computed in the 2D space defined by the projections of the population responses at the two time points.

#### Across-neurons correlations

For across-neuron correlations, we considered the same 100 randomly-defined pairs of neuronal pools that were used for encoding and consistency analysis (see “Analysis of stimulus encoding and consistency”). Multidimensional population responses of a given neuronal pool were first reduced to scalars by computing the projection along the direction of maximum stimulus variation in the space spanned by the population responses of the selected neuronal pool. Then, for each randomly-defined pair of disjoint pools, *γ* was computed in the 2D space defined by the projections of the population responses of the two pools.

### Mathematical model of encoding and readout with two neural features

We developed a simple two-features mathematical model of encoding and readout in a simulated stimulus discrimination task, to explore how possibly distinct roles of correlations in information encoding and information readout interact and impact task performance.

#### Neural encoding (stimulus-response) model

As far as the encoding model is concerned, we simulated the response of two generic features of neural activity (*r*_*1*_, *r*_*2*_) to two stimuli (*s = −1, s = 1*). Neural response distributions for a given stimulus are modelled as bivariate Gaussians *N*(***μ***_*s*_, **Σ**_s_), with mean and covariance matrix given by:

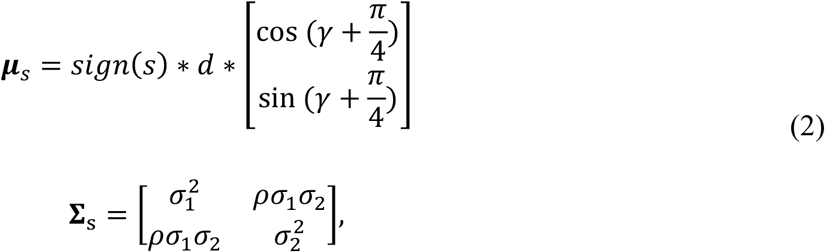

where 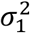 and 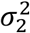 represent the variance of the neural responses along *r*_*1*_ and *r*_*2*_ respectively, while *ρ* determines the correlation between *r*_*1*_ and *r*_*2*_. For simplicity, in our simulations we chose *σ*_1_ = *σ*_2_ = *σ* and **Σ**_s1_ = **Σ**_s2_. We selected the means of the two response distributions (***μ***_*s*1_, ***μ***_*s*2_) to be symmetrically located around the origin in the 2D space defined by *r*_*1*_ and *r*_*2*_, at a distance *d*. Taken together, *d* and *σ* control the overlap between the two response distributions (in all our simulations we arbitrarily set 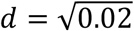 and *σ* = 0.3). The parameter *γ* controls the alignment between the signal correlations axis and the noise correlations axis, which are defined as the axis of maximum variability of trial-averaged responses at fixed stimulus and the axis of maximum trial-by-trial variability at fixed stimulus, averaged across stimuli, respectively. In our simulations, we varied *γ* by varying the signal correlations axis orientation (see Extended Data Fig. 2 and Supplementary Information): for *γ* = 0, ***μ***_*s*1_ and ***μ***_*s*2_ are located on the bisector of the first and third quadrants, resulting in perfect alignment between signal and noise; at increasing values of *γ*, the signal correlations axis direction is progressively rotated counterclockwise, resulting in more and more misalignment with the noise correlations axis; the maximum misalignment is reached when 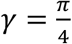 and signal and noise directions are orthogonal. We quantified the amount of stimulus information carried by the simulated neural responses as the fraction correct predictions of a linear decoder of stimulus identity applied to the simulated (*r*_*1*_, *r*_*2*_) responses. We further applied the same linear decoder to the responses along *r*_*1*_ and *r*_*2*_ separately for the computation of consistency. For the simulations in Fig. 2, Extended Data Fig. 2 and Extended Data Fig. 3 we performed, for a given choice of model parameters, 1000 simulations with 500 trials per stimulus.

#### Behavioral decision model

As far as the readout model is concerned, we simulated the process of generating a binary choice in each trial from neural activity through a logistic regression model:

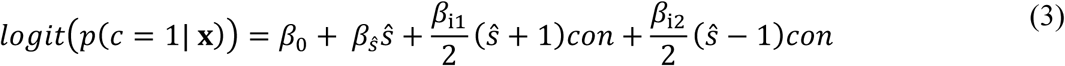

where:

- *ŝ* is a binary variable (*ŝ* = −1, *ŝ* = 1) which encodes the stimulus decoded from the concatenated activity of the two neural features;
- *con* is a binary variable that is 1 if the stimuli decoded individually from each neural feature are the same, and 0 otherwise;
- **x** indicates the entire set of predictors (*ŝ, con*).

The model coefficients *β*_0_, *β*_*ŝ*_, *β*_i1_ and *β*_i2_ control the relative impact of the different model predictors on the simulated choice. The values for the model coefficients were set as follows. We first defined a consistency modulation index *η*, ranging from 0 to 1, to control the relative strength of neural consistency in the readout. We then derived the readout efficacy, which we defined as the probability of conversion from *ŝ* to *c*, for each of the four possible combinations of predictors values, from the modulation index *η* and a reference readout efficacy *α*(*ŝ*) as:

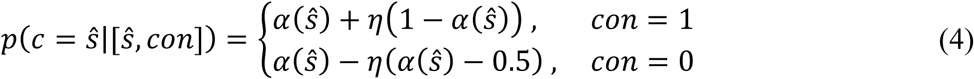

where *α*(*ŝ*) takes values between .5 and 1. For the simulations in Fig. 2, Extended Data Fig. 2 and Extended Data Fig. 3, we arbitrarily set *α*(0) = *α*(1) = 0.75. Given the readout efficacy values from equation (4), we used equation (3) to compute the model coefficients corresponding to the chosen modulation index *η*.

### Logistic regression of the mouse’s choice

To study the relevance of selected features of recorded neural population activity on the mouse’s choices, we fit to the recorded neural activity a logistic regression of the choice made by the animal in each trial (left/right turns). For each trial, we considered the choice *c* made by the mouse (*c =* left/right), the presented stimulus *s* (*s =* left/right), and the neural population activity for each pair of time points (across-time correlations analysis) or for each pair of randomly-selected neuronal pools (across-neuron correlations analysis) that were used for encoding and consistency analyses (see “Analysis of stimulus encoding and consistency”). For each session, trial split, and pair of time points or neuronal pools, we fitted the following logistic regression:

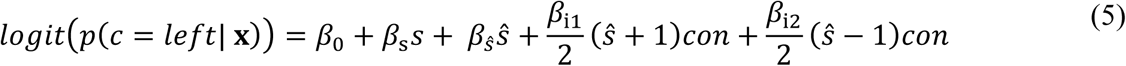

where

- *ŝ* is a binary variable (*ŝ =* left/right) that encodes the most likely stimulus decoded from the concatenated activity of two time points or neuronal pools;
- *con* is a binary variable that is 1 if the stimuli decoded individually from each time point or neuronal pool are the same, and 0 otherwise;
- **x** indicates the entire set of neural (*ŝ, con*) and non-neural (*s*) predictors.

Logistic regression fitting was implemented with custom Python software, by building on the *statsmodel* module^53^. Note that we used the convention (left = 1, right = −1) to map the categorical variables *s, ŝ, c* to numerical variables suitable for logistic fitting. For each trial split defined for encoding and consistency analyses (see “Analysis of stimulus encoding and consistency”), the logistic regression was fit on trials belonging to the testing set, for which there exists a prediction of the most likely stimulus from population activity for each individual and pair of time points or neuronal pools. The logistic regression was fit using L1-regularized maximum likelihood, for a conservative selection of the predictors of the mouse’s choice. The regularization hyperparameter (*λ*) was selected by maximizing 3-fold cross-validated fraction of deviance explained (FDE, see “Predictive performance and cross validation”). The few instances where the maximum-likelihood optimization did not converge numerically were not considered for further analyses.

For control analyses, we fitted mouse’s choices with more complex choice regressions that included other predictors on top of those described in equation (5).

In order to discern the genuine role of across-time neural consistency (*con*) in explaining the mouse’s choices from that of across-time behavioral consistency (*con*_*b*_), we fitted a logistic regression which included additional behavioral consistency-dependent predictors:

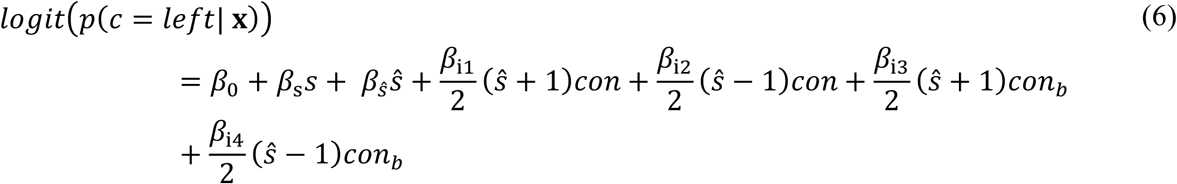

We performed this control analysis for three behavioral parameters of interest that were measured during the experiments: the lateral running velocity, lateral position and view angle of the mouse in the virtual environment (see Extended Data Fig. 5 and Supplementary Information). Two values of lateral running velocity or lateral position taken at two different time points were defined to be consistent whenever their sign was the same; two values of view angle at two different time points were defined to be consistent whenever they were both higher or both lower than 90°.

In order to resolve the genuine role of across-time or across-neurons consistency in explaining the mouse’s choices, above and beyond the overall level of stimulus information encoded in neural activity, we fitted mouse’s choices with a logistic regression where the discrete binary variable *ŝ* in equation (5) was replaced with the corresponding continuous value of the decoder stimulus posterior probability *p*(*s* = *left*|***r***) (see “Analysis of stimulus encoding and consistency”). This choice regression, differently from the one of equation (5), also accounts for the confidence associated to a given stimulus decoded from the concatenated activity of two time points or two neuronal pools.

#### Predictive performance and cross validation

To test for the significance of the contribution of a predictor (or set of predictors) in explaining the mouse’s choices, we compared the predictive performance of the regression fitted (and tested) on predictors values extracted from the recorded data with the predictive performance of a regression fitted (and tested) on artificial data in which the values of the predictor (or set of predictors) of interest were shuffled across trials. Predictive performance was quantified as the cross-validated fraction of deviance explained (FDE). FDE was evaluated with 3-fold cross validation, thus addressing concerns of overfitting. For each fold, we computed the log-likelihood *l* of the test data given the values of the *β* coefficients of the regression fit on the training data. To calculate a reference null value for the log-likelihood, we computed the log-likelihood *l*_0_ of the test data given the value of the coefficient *β*_0_ of an intercept-only regression fit on the training data. The FDE was then defined as:

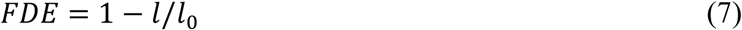

### Estimating the impact of across-time and across-neuron correlations on task performance

To estimate the impact of across-time and across-neuron correlations on mouse’s task performance, we generated synthetic choices using the experimentally-fit choice regression of equation (5). As input to the regression, we provided either predictors extracted from the real recorded neural data, which included across-time and across-neuron correlations, or predictors extracted from hypothetical neural data where correlations of interest were artificially removed by shuffling (see “Removing across-time or across-neuron noise correlations by shuffling”). We used this analytic approach because current experimental methods cannot remove correlations online during task performance, and thus we had to estimate effects with post-hoc removal of correlations.

Technically, given the discrete nature of the predictors used in the choice regression of equation (5), we did not need to generate synthetic choices for the computation of the task performance *p*(*c* = *s*), which is the probability that the choice *c* matches the presented stimulus *s*. Rather, we simply computed the probabilities *p*(**X**) for all possible combination of predictors values X, we multiplied them with the corresponding readout probabilities *p*(*c* = *s*| **X**) obtained from the logistic choice regression, and then summed over all possible combinations of predictors values X as follows (see Extended Data Fig. 1 for a schematic representation of this operation):

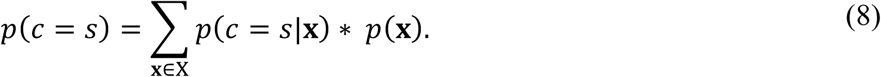

Note that the same readout probabilities *p*(*c* = *s*|**X**) were used for the computation of task performance from both real and shuffled neural data.

We further isolated the part of task performance that can be attributed to the (real or shuffled) neural activity, by subtracting from the total estimated task performance a baseline non-neural estimated task performance. The non-neural task performance was computed by generating choices (or applying equation (8)) after shuffling the values of neural predictors (i.e. predictors defined from features of neural activity) across trials, while keeping the relationship between non-neural predictors (i.e. predictors that did not depend on neural activity) and mouse’s choices fixed. Note that the shuffling procedure allowed to discard uninteresting differences between task performance from real neural data and that from shuffled neural data that are due to intrinsic differences between the marginal distributions of the neural predictors. The estimated value of task performance due to the recorded neurons is by definition non-negative and it is identically zero when the coefficients associated to neural predictors are zero, as we intuitively require to a measure of the behavioral impact of the neural predictors.

### Computation of readout efficacy of the transformation from stimulus information to choice

We termed “readout efficacy” *p*(*c* = *ŝ*) the probability of conversion from stimulus encoded in neural activity to choice associated to a given readout. Given the discrete nature of the predictors used in the choice regression of equation (5), similarly to what we did for the estimation of task performance (see “Estimating the impact of across-time and across-neuron correlations on task performance”), we computed the efficacy of the enhanced-by-consistency readout as:

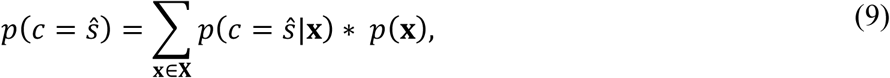

where *X* represents the set of all possible combinations of predictors values and *p*(*c* = *ŝ*|**X**) are obtained from the logistic choice regression.

To generate the readout maps in Fig. 3c, f we computed, separately for consistent and inconsistent trials, readout efficacy as a deviation from a stimulus-driven baseline, which is the average probability of choice being left or right when the presented stimulus is left or right, as:

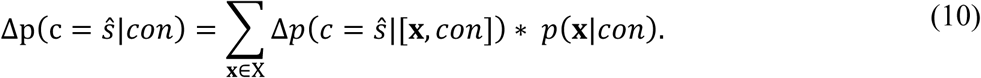

In equation (10), *X* represents the set of all possible combinations of [*s, ŝ*] predictors values and *Δp*(*c* = *ŝ*|[**x**, *con*]) = *p*(*c* = *ŝ*|[*s, ŝ, con*]) − *p*(*c* = *ŝ*|*s*).

### Matching enhanced-by-consistency and consistency-independent readouts in terms of efficacy

To quantify the impact on task performance of the enhanced-by-consistency experimentally-measured readout, we compared the task performance predicted by the experimentally-fit choice regression to the one predicted by a consistency-independent choice regression of the form:

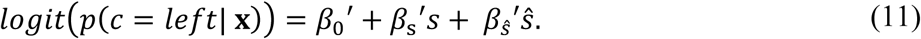

For a fair comparison, the values of the coefficients 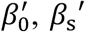 and 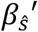 were chosen so that the two readouts were matched in terms of readout efficacy (see “Computation of readout efficacy in the transformation from stimulus information to choice”). Note how the readout efficacy *p*(c = *ŝ*) in equation (9) depends on both the readout probabilities *p*(*c* = *ŝ*|**x**) and the predictors probabilities *p*(**x**). Thus, the condition of matched readout efficacy is specific for a given input to the regressions.

Concretely, in order to compute the values of the coefficient of the consistency-independent choice regression in (11), we imposed the following conditions:

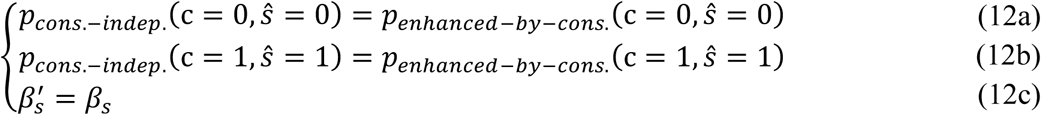

then appropriately plugged equations (5) and (11) into (12a) and (12b), and solved for 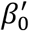 and 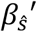.

### Removing across-time or across-neuron noise correlations by shuffling

To study the role of across-time and across-neuron correlations on stimulus encoding and task performance, we generated artificial neural population vectors in which we removed correlations by a shuffling procedure.

#### Across-time correlations analysis

Across-time correlations between population vectors at different time points were removed by shuffling trial identities independently for each of the two population vectors within subsets of trials of the same stimulus category. This procedure ensured that “across-time signal correlations” were maintained, while “across-time noise correlations” were disrupted. Note that single-cell autocorrelations were also disrupted with this shuffling procedure.

#### Across-neuron correlations analysis

Across-neuron correlations between two neuronal pools were removed by shuffling trial identities independently for each pool within subsets of trials of the same stimulus category. This procedure ensured that “across-neuron signal correlations” were maintained, while “across-neuron noise correlations” were disrupted. Note that classical signal correlations between any pairs of neurons from the two disjoint pools were thus maintained, while classical noise correlations between any pairs of neurons from the two disjoint pools were disrupted.

Shuffling was performed separately within trials of the training and the testing set.

### Data and code availability

The data and code that support the findings of this study are available from the corresponding authors upon reasonable request.

## References

1. Ni, A. M., Ruff, D. A., Alberts, J. J., Symmonds, J. & Cohen, M. R. Learning and attention reveal a general relationship between population activity and behavior. Science 359, 463–465 (2018).

2. Kohn, A., Coen-Cagli, R., Kanitscheider, I. & Pouget, A. Correlations and Neuronal Population Information. Annu. Rev. Neurosci. 39, 237–256 (2016).

3. Panzeri, S., Harvey, C. D., Piasini, E., Latham, P. E. & Fellin, T. Cracking the Neural Code for Sensory Perception by Combining Statistics, Intervention, and Behavior. Neuron 93, 491–507 (2017).

4. Gawne, T. J. & Richmond, B. J. How Independent Are the Messages Temporal Cortical Neurons? by Adjacent Inferior. The Journal of Neuroscience 73, (1993).

5. Averbeck, B. B., Latham, P. E. & Pouget, A. Neural correlations, population coding and computation. Nature Reviews Neuroscience 7, 358–366 (2006).

6. Zohary, E., Shadlen, M. N. & Newsome, W. T. Correlated neuronal discharge rate and its implications for psychophysical performance. Nature 370, 140–143 (1994).

7. Moreno-Bote, R. et al. Information-limiting correlations. Nat. Neurosci. 17, 1410–1417 (2014).

8. Bartolo, R., Saunders, R. C., Mitz, A. R. & Averbeck, B. B. Information limiting correlations in large neural populations. J. Neurosci. 2072–19 (2020).

9. Rumyantsev, O. I. et al. Fundamental bounds on the fidelity of sensory cortical coding. Nature 1–6 (2020). doi:10.1038/s41586-020-2130-2

10. Diesmann, M., Gewaltig, M. O. & Aertsen, A. Stable propagation of synchronous spiking in cortical neural networks. Nature 402, 529–533 (1999).

11. Zandvakili, A. & Kohn, A. Coordinated Neuronal Activity Enhances Corticocortical Communication. Neuron 87, 827–839 (2015).

12. Salinas, E. & Sejnowski, T. J. Correlated neuronal activity and the flow of neural information. Nature Reviews Neuroscience 2, 539–550 (2001).

13. Zylberberg, J., Pouget, A., Latham, P. E. & Shea-Brown, E. Robust information propagation through noisy neural circuits. PLoS Computational Biology 13(4), e1005497 (2017).

14. Abeles, M. Corticonics : neural circuits of the cerebral cortex. (Cambridge University Press, 1991).

15. Harvey, C. D., Coen, P. & Tank, D. W. Choice-specific sequences in parietal cortex during a virtual-navigation decision task. Nature 484, 62–68 (2012).

16. Minderer, M., Brown, K. D. & Harvey, C. D. The Spatial Structure of Neural Encoding in Mouse Posterior Cortex during Navigation. Neuron 102, 232-248.e11 (2019).

17. Runyan, C. A., Piasini, E., Panzeri, S. & Harvey, C. D. Distinct timescales of population coding across cortex. Nature 548, 92–96 (2017). doi:10.1038/nature23020

18. Hanks, T. D. et al. Distinct relationships of parietal and prefrontal cortices to evidence accumulation. Nature 520, 220–223 (2015).

19. Morcos, A. S. & Harvey, C. D. History-dependent variability in population dynamics during evidence accumulation in cortex. Nat. Neurosci. 19, 1672–1681 (2016).

20. Raposo, D., Kaufman, M. T. & Churchland, A. K. A category-free neural population supports evolving demands during decision-making. Nat. Neurosci. 17, 1784–1792 (2014).

21. Pho, G. N., Goard, M. J., Woodson, J., Crawford, B. & Sur, M. Task-dependent representations of stimulus and choice in mouse parietal cortex. Nat. Commun. 9, 2596 (2018).

22. Goard, M. J., Pho, G. N., Woodson, J. & Sur, M. Distinct roles of visual, parietal, and frontal motor cortices in memory-guided sensorimotor decisions. Elife 5, (2016).

23. Akrami, A., Kopec, C. D., Diamond, M. E. & Brody, C. D. Posterior parietal cortex represents sensory history and mediates its effects on behaviour. Nature 554, 368–372 (2018).

24. Averbeck, B. B. & Lee, D. Effects of noise correlations on information encoding and decoding. J. Neurophysiol. 95, 3633–3644 (2006).

25. Nogueira, R. et al. The effects of population tuning and trial-by-trial variability on information encoding and behavior. J. Neurosci. 40, 1066–1083 (2020).

26. Romo, R., Hernández, A., Zainos, A. & Salinas, E. Correlated neuronal discharges that increase coding efficiency during perceptual discrimination. Neuron 38, 649–657 (2003).

27. Reich, D. S., Mechler, F. & Victor, J. D. Independent and redundant information in nearby cortical neurons. Science 294, 2566–2568 (2001).

28. Gold, J. I. & Shadlen, M. N. Neural computations that underlie decisions about sensory stimuli. Trends Cogn. Sci. 5, 10–16 (2001).

29. Kanitscheider, I., Coen-Cagli, R. & Pouget, A. Origin of information-limiting noise correlations. Proc. Natl. Acad. Sci. U. S. A. 112, E6973–E6982 (2015).

30. Zariwala, H. A., Kepecs, A., Uchida, N., Hirokawa, J. & Mainen, Z. F. The limits of deliberation in a perceptual decision task. Neuron 78, 339–351 (2013).

31. Mazurek, M. E. & Shadlen, M. N. Limits to the temporal fidelity of cortical spike rate signals. Nat. Neurosci. 5, 463–471 (2002).

32. Renart, A. et al. The asynchronous state in cortical circuits. Science 327, 587–590 (2010).

33. Ecker, A. S. et al. Decorrelated neuronal firing in cortical microcircuits. Science 327, 584–587 (2010).

34. Shahidi, N., Andrei, A. R., Hu, M. & Dragoi, V. High-order coordination of cortical spiking activity modulates perceptual accuracy. Nat. Neurosci. 22, 1148–1158 (2019).

35. Histed, M. H. & Maunsell, J. H. R. Cortical neural populations can guide behavior by integrating inputs linearly, independent of synchrony. Proc. Natl. Acad. Sci. U. S. A. 111, E178–E187 (2014).

36. Francis, N. A. et al. Small Networks Encode Decision-Making in Primary Auditory Cortex. Neuron 97, 885–897 (2018).

37. Emiliani, V., Cohen, A. E., Deisseroth, K. & Häusser, M. All-optical interrogation of neural circuits. Journal of Neuroscience 35, 13917–13926 (2015).

38. Stringer, C., Michaelos, M. & Pachitariu, M. High precision coding in visual cortex. bioRxiv 679324 (2019). doi:10.1101/679324

39. O’Connor, D. H. et al. Neural coding during active somatosensation revealed using illusory touch. Nat. Neurosci. 16, 958–965 (2013).

40. Pitkow, X., Liu, S., Angelaki, D. E., DeAngelis, G. C. & Pouget, A. How Can Single Sensory Neurons Predict Behavior? Neuron 87, 411–423 (2015).

41. Nirenberg, S., Carcieri, S. M., Jacobs, A. L. & Latham, P. E. Retinal ganglion cells act largely as independent encoders. Nature 411, 698–701 (2001).

42. Karpas, E., Maoz, O., Kiani, R. & Schneidman, E. Strongly correlated spatiotemporal encoding and simple decoding in the prefrontal cortex. bioRxiv 693192 (2019). doi:10.1101/693192

43. Berens, P. CircStat : A MATLAB Toolbox for Circular Statistics. J. Stat. Softw. 31, 1–21 (2009).

## Methods references

44. Harvey, C. D., Collman, F., Dombeck, D. A. & Tank, D. W. Intracellular dynamics of hippocampal place cells during virtual navigation. Nature 461, 941–946 (2009).

45. Aronov, D. & Tank, D. W. Engagement of Neural Circuits Underlying 2D Spatial Navigation in a Rodent Virtual Reality System. Neuron 84(2), 442–456 (2014). doi:10.1016/j.neuron.2014.08.042

46. Greenberg, D. S. & Kerr, J. N. D. Automated correction of fast motion artifacts for two-photon imaging of awake animals. J. Neurosci. Methods (2009). doi:10.1016/j.jneumeth.2008.08.020

47. Shi, J. & Malik, J. Normalized cuts and image segmentation. IEEE Trans. Pattern Anal. Mach. Intell. (2000). doi:10.1109/34.868688

48. Vogelstein, J. T. et al. Fast nonnegative deconvolution for spike train inference from population calcium imaging. J. Neurophysiol. 104, 3691–3704 (2010).

49. Boser, B. E., Guyon, I. M. & Vapnik, V. N. A training algorithm for optimal margin classifiers. in Proceedings of the fifth annual workshop on Computational learning theory - COLT ‘92 144–152 (ACM Press, 1992). doi:10.1145/130385.130401

50. Cortes, C. & Vapnik, V. Support-vector networks. Mach. Learn. 20, 273–297 (1995).

51. Chang, C.-C. & Lin, C.-J. LIBSVM: A Library for Support Vector Machines. (2001).

52. Lin, H.-T., Lin, C.-J. & Weng, R. C. A note on Platt’s probabilistic outputs for support vector machines. Mach. Learn. 68, 267–276 (2007).

53. Seabold, S. & Perktold, J. Statsmodels: Econometric and Statistical Modeling with Python. PROC. OF THE 9th PYTHON IN SCIENCE CONF (2010).

