## Supplementary Information for "Correlations enhance the behavioral readout of neural population activity in association cortex"

#### S.1 Systematic analysis of the predictions of the two-feature encoding-readout model

In Fig. 2 and in the main text we introduced a simple mathematical model of how two generic features of neural activity encode information about a binary stimulus, and how this information is read out to inform choice in a simulated stimulus discrimination task. We introduced a parametric model of readout that expresses the probability of simulated choices as a function of both the identity of the stimulus encoded by the two features jointly and the consistency of the stimulus encoded by each of the two individual features separately. This readout model has a parameter, indicated by $\eta$ and termed consistency modulation in the readout, that expresses how much the behavioral choice is modulated, for a given stimulus encoded in joint population activity, by whether or not stimulus information is consistent across the two neural features. Positive values of $\eta$ correspond to an enhanced-by-consistency readout, in which behavioral choice is more likely to follow the stimulus encoded in neural activity when stimulus information is consistent. A zero value of $\eta$ corresponds to the traditionally-considered consistency-independent readout, in which the probability of choice following the stimulus encoded in neural activity is independent of stimulus information consistency. The two-features encoding and readout model is defined mathematically in Methods, Section "Mathematical model of encoding and readout with two neural features".

Here, we investigate the conditions in which an enhanced-by-consistency readout overcomes the information-limiting nature of correlations and provides a benefit for task performance (as in Fig. 2k), by varying model parameters. In brief, our simulations show that the benefit for task performance provided by the enhanced-by-consistency readout in the presence of information limiting-correlations comes about because this readout uses more efficiently the stimulus information to inform choices when neural population activity is consistent, which is more likely in the presence than in the absence of correlations. In particular, a performance benefit is present when correlations boost the overall readout efficiency more than they damage information encoding.

To better understand the conditions in which the enhanced-by-consistency readout can overturn the information deficit due to correlations, we studied systematically how the effect of correlations on information coding and task performance depended on key model parameters: the angle $\gamma$ between the signal and noise correlation axes, the strength $\eta$ of the consistency modulation in the readout, and the strength $\rho$ of noise correlations (all parameters are defined in Methods, Section "Mathematical model of encoding and readout with two neural features"). Results are reported in Extended Data Fig. 2 and were briefly summarized in the main text. Here we report them in more detail, highlighting some more points of interest arising from the analysis.

We first fixed an arbitrary value of $\eta = 0.7$, corresponding to an enhanced-by-consistency readout, and considered the effect of the misalignment angle $\gamma$ between signal and noise correlations axes (Extended Data Fig. 2b-e). Three different examples of stimulus-specific response distributions along the two features $r_1$, $r_2$ for three different values of $\gamma$ (two values corresponding to information-limiting correlations, and one value corresponding to information-enhancing correlations) are reported in Extended Data Fig. 2a. In agreement with previous work[1–4], we found that increasing the misalignment $\gamma$ progressively decreased (and for higher $\gamma$ values, even overturned) the stimulus information loss of correlated responses over uncorrelated responses (Extended Data Fig. 2b).

It is interesting to consider the range of $\gamma$ values in which, for this model, information is decreased by correlations but task performance is enhanced by it. This range is highlighted as a red dashed rectangle in Extended Data Fig. 2b-e. The impact of information-limiting correlations on task performance, under the hypothesis of an enhanced-by-consistency readout, results from a tradeoff between the increase of decoding errors due to correlations and the increase of consistency due to correlations. The first effect negatively impacts performance, whereas the second one might contribute positively or negatively, depending on the relationship between consistency and correct decoding. Indeed, the enhanced-by-consistency readout amplifies both the faithful stimulus information present in correctly decoded trials that are consistent and the misleading stimulus information present in incorrectly decoded trials that are consistent. The net effect of correlations on task performance depends on the tradeoff of these two effects. Increasing $\gamma$ increases the fraction of correctly decoded trials that are consistent in correlated data and decreases this fraction in uncorrelated data (Extended Data Fig. 2c). Increasing $\gamma$ also decreases the fraction of incorrectly decoded trials that are consistent in correlated data but not in shuffled data (Extended Data Fig. 2d). Both effects contribute to increase the task performance produced by the enhanced-by-consistency readout when $\gamma$ increases. The combined effect of a reduction of the loss of information due to correlations, together with the increase in the proportion of trials with correct decoding that are more efficiently readout because they are consistent, together results in a range of $\gamma$ values for which noise correlations are, at the same time, information limiting and task performance enhancing (Extended Data Fig. 2e). Although the parameter values of multidimensional neural data may not be straightforwardly comparable to those of our simple two-feature model, it is interesting to note that our two-feature model predicts that the enhanced-by-consistency readout can overturn the information deficit of correlations in a range of non-zero but small values of $\gamma$ that roughly matches the range of $\gamma$ values measured in on our experimental data (Fig. 1e, j).

Another interesting finding regarding the dependence of the task performance on $\gamma$ is that higher task performance with correlated data is found also for higher values of $\gamma$ in which correlations enhance, rather than limit, information in neural activity (compare Extended Data Fig. 2b with Extended Data Fig. 2e). This result suggests that while the enhanced-by-consistency readout is particularly good at overturning, at the behavioral level, the information deficit of information-limiting, it also preserves, at the behavioral level, their information advantage when they are information-enhancing.

To study how the range of information limiting, task performance enhancing $\gamma$ values depends on the strength by which consistency affects readout, we systematically varied in our simulations the consistency modulation index $\eta$. We found (Extended Data Fig. 2f, g, top) that the stronger the consistency modulation in the readout, the larger the range of information limiting, task performance enhancing $\gamma$ values. Importantly, when consistency did not modulate the readout much (small values of $\eta$), there was no advantage in task performance when correlations were information limiting (Extended Data Fig. 2 f, g, top), and correlations were stronger in error than in correct trials (Extended Data Fig. 2 h, top).

Increasing the strength of noise correlations among features (Extended Data Fig. 2f,g, bottom) also affected the relationship between encoded information and task performance, with the above described pattern of task performance advantage of correlation found for all levels of correlations, but with weaker correlations resulting in information limiting, task performance enhancing cases happening at higher $\gamma$ values.

If the readout was independent of consistency (case $\eta = 0$; corresponding to the values at the top of the upper panels of Extended Data Fig. 2f,g), as expected by simple intuition, performance in the task would follow directly the level of information encoded in population activity (the higher the information the higher the task performance). In this case, when correlations are information limiting (lower $\gamma$ values) then task performance is higher in shuffled data, and when correlations are information enhancing (higher $\gamma$ values) then task performance is higher in correlated data. Note that, with information-limiting correlations and consistency-independent readout, correlations tend to be stronger in error than in correct trials (Extended Data Fig. 2h).

We further observed that the difference in the measured correlation strength between trials with correct and error choices, when the readout is enhanced by consistency, shows a pattern of dependence on the model parameters that resembles, but does not match, the advantage of correlated over uncorrelated data in task performance (compare Extended Data Fig. 2g with Extended Data Fig. 2h). In particular, in our model we found some conditions in which information-limiting noise correlations are stronger in correct trials, without them having a positive effect on task performance. However, as we reported above, this did not happen if the readout was consistency-independent, in which case correlations were higher in error trials when they limited both information and task performance. Thus, we conclude that, according to our simple model, observing higher correlations in correct trials when correlations are information-limiting (as we found in PPC data, see Fig. 1) is indicative that consistency may affect readout, but it is not sufficient to imply an advantage of correlations in task performance. The latter question should be instead investigated by comparing task performance resulting from the readout of correlated data with the one resulting from the readout of uncorrelated data.

**S.2 Using the two-features encoding-readout model to compare the task performance predicted by enhanced-by-consistency and consistency-independent readouts.**

In this section, we show that our two-feature encoding-readout model predicts that the enhanced-by-consistency readout provides and advantage for task performance when compared to a consistency-independent readout, in case neural responses are characterized by information-limiting correlations.

For a given set of parameters of the encoding model, describing how the two correlated features encode stimulus information, we computed and compared simulated task performance obtained with either an enhanced-by-consistency readout (with consistency modulation index $\eta$) or with a consistency-independent readout (consistency modulation index $\eta = 0$). When comparing the two readouts, we set the logistic regression coefficients such that the two readouts were matched in terms of readout efficacy, which is the probability of transformation from stimulus encoded in population activity into choice (see Methods, Section "Matching enhanced-by-consistency and consistency-independent readouts in terms of efficacy"). In other words, the two readouts are matched in terms of fraction of times the choice predicted by the readout model matches the stimulus decoded from neural activity. The difference is that in the consistency-independent readout this fraction is uniform across trial types, whereas in the enhanced-by-consistency readout this fraction is higher in consistent than in inconsistent trials.

Results are reported in Extended Data Fig. 3 for a range of values of the signal-noise angle $\gamma$, the consistency parameter $\eta$, the strength of noise correlation $\rho$. The results show that, in general, when information is encoded in the presence of information-limiting correlations (low values of the signal-noise angle $\gamma$), having an enhanced-by-consistency readout is advantageous for task performance. The behavioral advantage is maximal when $\gamma = 0$ and decreases as $\gamma$ increases (Extended Data Fig. 3a). As expected, the difference in task performance predicted by the two readout models is larger when the strength of the consistency modulation is larger (Extended Data Fig. 3b).

**S.3 The effect of neural consistency on the mouse's single trial choices cannot be explained by the fact that consistent trials may also have more overall stimulus information**

In Fig. 3 we showed that the mouse's choice is more likely to follow the stimulus encoded in PPC neural activity in trials in which neural information is consistent across time or across neurons, than in trials in which neural information is inconsistent. This finding suggests that the readout mechanism performs more efficiently in the presence of consistent neural representations.

A possible concern with these analyses is that the differences in mouse's task performance between consistent and inconsistent trials might be due to higher stimulus information in neural activity in consistent trials. That consistent trials may have on average higher information than inconsistent trials is intuitive, and it is illustrated with a cartoon in Extended Data Fig. 4a, which depicts the distribution of responses of two neural features to two different stimuli. As in Fig. 2, the black line indicates the linear decoding boundary that best separates the responses to the two stimuli. The colored background represents, for each combination of values of the features $r_1$ $r_2$, the corresponding value of the decoder posterior probability $p(s=1|\mathbf{r})$ that stimulus s =1 has occurred given the observation of neural response **r**. From this figure, it is apparent that the posterior probability $p(s=1|\mathbf{r})$ is on average more different from 0.5, at thus more "informative" about stimulus s = 1 or stimulus s = -1, in regions of the $r_1$, $r_2$ space corresponding to consistent trials than in regions corresponding to inconsistent trials.

To address this concern, we repeated the analyses of Fig. 3 on trials partitioned into those with low, medium, or high "stimulus information" (Extended Data Fig. 4). In more detail, we partitioned trials into three groups according to how much the stimulus posterior differed from 0.5: low: < 0.16; medium: 0.16-0.32; or high: > 0.32. In both of our datasets, mouse's task performance in correctly (respectively incorrectly) decoded trials with the same or even with lower stimulus information level, was still higher (respectively lower) when information was consistent across neurons or across time (Extended Data Fig. 4c-e and g-i).

To further control for potential confounders due to differences in the levels of stimulus information between trials with consistent and inconsistent stimulus information, we fitted to the data a more refined logistic regression that used the posterior probability of the stimulus given the observed neural responses as predictor of the mouse's choices, rather than just the identity of the most likely stimulus given the neural responses. In more detail, the refined regression was defined by replacing, in the regression discussed in the main text, the binary stimulus $\hat{s}$ decoded from PPC activity with the corresponding continuous posterior probability $p(s=1|\mathbf{r})$ (see Methods). We found that, in both datasets, consistency provided a similar contribution to predicting choices when we used the more sophisticated logistic regression in which we included the magnitude of the stimulus information, instead of only the identity of the decoded stimulus (Extended Data Fig. 4b, f).

Therefore, our finding that information in neural activity informs choice more effectively when it is consistent cannot be explained by differences in overall stimulus information level. Rather, these results suggest that, for a given amount of sensory information, more information can be extracted to guide behavioral choices if it is distributed redundantly across neurons or across time.

**S.4 The role of neural consistency in the readout of PPC activity is not due to the consistency of measured behavioral parameters**

In Fig. 3 we used choice regressions of neural population activity to show that, in the sound localization task PPC dataset, the across-time consistency of stimulus information encoded in neural population activity impacts behavioral choices.

Here, we performed control analyses to rule out the concern that the impact on the mouse's choice of across-time consistency of PPC activity does not only reflect the effect of some features of the animal's running, whose temporal consistency may correlate with both the mouse's choice and the temporal consistency of neural activity.

To control for potential contributions from running-related neural activity to the consistency-dependent terms of our experimentally-fit choice regression, we developed and fit to PPC data a more sophisticated choice regression that explicitly includes such contributions. In more detail, we developed an extended version of the choice regression presented in the main text, in which we explicitly included behavioral consistency as an additional predictor of the animal's choice (see Methods, Section "Logistic regression of the mouse's choice"). We first defined the across-time behavioral consistency for three different behavioral parameters that were recorded during the experiments: the mouse lateral running velocity, lateral position, and view angle in the virtual maze (see Extended Data Fig. 5a for some example traces). We then fit to the PPC neural population data of the sound localization task dataset a logistic regression model of the choice of the animal that included as predictors, in addition to the presented stimulus, the decoded stimulus, and the two neural consistency interaction terms, also two behavioral consistency interaction terms (see Methods, Section "Logistic regression of the mouse's choice").

We found that across-time neural consistency improved choice predictability above and beyond that obtained by considering across-time behavioral consistency only, for all three considered behavioral parameters (Extended Data Fig. 5c-e). This was established by comparing the fraction of deviance explained by the extended choice regression including neural and behavioral consistency-dependent predictors with the fraction of deviance explained by the same type of regression but fitted after shuffling neural consistency-dependent predictors across trials.

The fact that neural consistency still contributed to predicting choices when we added the consistency of running-related variables to the choice regression suggests that consistency of the instantaneous PPC population activity across time genuinely influences the behavioral readout of the stimulus information, above and beyond what can be predicted about choice from the consistency of measured behavioral variables.

### S.5 Comparisons of the effect of across-time correlations in auditory cortex and posterior parietal cortex in the sound localization task

We repeated all the analyses of the role of across-time correlations, that we described in Fig. 3-4 for the PPC neural recordings acquired during the sound localization task, on the auditory cortex (AC) neural population recordings available for the same task.

Previous analyses of these data reported a far lower strength of noise correlations in AC, relative to PPC, suggesting that sensory and association areas may differ in their levels of correlations[5]. Here, we focused on estimating the effects of such noise correlations on stimulus coding and behavioral readout.

With regard to stimulus coding in AC, we observed overall a higher stimulus decoding performance in AC than in PPC (Extended Data Fig. 6a, g), compatible with the view that AC is an area strongly involved in the encoding of sound information. We found that across-time correlations had an information limiting effect and reduced consistency of information also in AC; however, both such effects of across-time correlations were far smaller in AC than in PPC (Extended Data Fig. 6a, b, g, h).

With regard to the estimation of the dependence on consistency of the behavioral readout of AC activity, we found that across-time consistency of stimulus information in AC provided negligible improvements in behavioral choice predictions when compared to the effect of across-time consistency in PPC (much smaller increase in cross-validated Fraction of Deviance Explained in AC than in PPC when comparing the full choice regression including consistency with the “No Cons” regression that is fit on shuffled consistency-dependent predictors, see Extended Data Fig. 6c, i, and smaller values of the coefficients for the consistency-dependent interaction terms in AC than in PPC, see Extended Data Fig. 6d, j).

With regard to the estimated effect of across-time correlations on task performance, we found that, in contrast with our findings in PPC (Extended Data Fig. 6e), only a small and weakly significant increase of task performance was predicted with correlated data in AC (Extended Data Fig. 6k). Finally, and still as a consequence of the small effect of correlations on both encoding and readout in AC data, task performance from the readout of AC data would be expected to not differ significantly with either a consistency-independent-readout or with the best-fit readout (Extended Data Fig. 6l).

### S.6 Comparisons of the effect of across-neuron correlations in auditory cortex and posterior parietal cortex in the sound localization task

We repeated all the analyses concerning across-neuron correlations, that we described in Fig.3-4 for the PPC neural recordings acquired during the visual evidence accumulation task, on both the AC and the PPC neural recordings acquired during the sound localization task.

With regard to stimulus coding in PPC, we observed (Extended Data Fig. 7a, g) overall a lower decoding performance from PPC data in the sound localization dataset than in the evidence accumulation dataset, because of the lower number of neurons per session recoded in the sound localization dataset (~50 vs ~350, see Methods).

With regard to the estimation of the dependence on across-neuron consistency of the behavioral readout of PPC activity, we found that all the results found in the PPC evidence accumulation dataset were confirmed in the PPC sound localization dataset. In particular, in the PPC sound localization dataset, consistency due to across-neuron correlations significantly increased the prediction of choice from PPC activity (Extended Data Fig. 7c, d). Importantly, our analysis estimated higher task performance in the sound localization task for PPC activity with across-neuron correlations, with respect to shuffled PPC data (Extended Data Fig. 7e). Our analyses also estimated that having an enhanced-by-consistency readout would be beneficial for task performance when considering across-neuron noise correlations in PPC, with respect to having a readout that is independent of consistency (Extended Data Fig. 7f).

With regard to the AC recordings, we found (Extended Data Fig. 7g-l) that the effect of across-neuron correlations on stimulus encoding, behavioral readout, and estimated task performance was very small, similar to what we reported in Extended Data Fig. 6g-l for the role of across-time correlations.

Overall, these results confirm that the beneficial effect for behavioral readout of across-neuron noise correlations in association cortices holds both for the sound localization and the visual evidence accumulation task.

**S.7 Evidence supporting a non-optimal readout of stimulus information in population activity for perceptual discrimination performance**

Our conceptual distinction between consistency-independent and enhanced-by-consistency readout models implicitly assumes that there is some suboptimality in the behavioral readout of stimulus information encoded in neural activity for task performance. More specifically, it assumes that the animal's choice does not always follow the stimulus decoded from neural activity, because not all stimulus information present in neural activity is used for behavior. This suboptimality assumption is critical for our study, because if we assumed choice to always follow the stimulus decoded from neural activity, then both readout models would operate exactly in the same way on the population activity and predict the same behavioral decision in all trials.

In this Section we thus consider whether our results support the idea that not all stimulus information carried by neural activity is used toward producing a correct behavioral choice, or in other words that the stimulus information in population activity is not read out optimally to form behavioral decisions. Operationally, a first possible definition of optimal readout is a readout that uses for perceptual discrimination behaviors all stimulus information in neural activity. Verifying this hypothesis requires comparing the amount of stimulus information carried by neural population activity with the amount stimulus information in neural activity that is used for behavioral choice. Suppose that stimulus information in the whole brain is made of the "recorded information" contained in the neurons under consideration (and perhaps carried in a redundant way by other non-recorded neurons) and by the "non-recorded information" carried by the non-recorded neurons and non-redundant with the information carried by the recorded neurons. In our choice regression used to estimate the readout of real neural population data, we included the effect of these two sources of information on behavioral choices. The effect of the recorded information is accounted for by what we indicated as neural predictors, i.e. the decoded stimulus predictor and the two consistency-dependent predictors. The effect of the non-recorded information is instead incorporated by the stimulus predictor, which includes the effect of the stimulus information carried by the non-recorded neurons, above and beyond that carried by the recorded neurons. (In addition, we also included a bias term that includes the overall effect of non-stimulus-informative contributions of neural activity). If all stimulus information in the recorded neural activity is read out optimally, we expect that the predicted contribution of the recorded population to task performance (estimated from real data with the choice regression) is equal to the stimulus information that is present in neural activity. In our data, we quantified stimulus information contained in the recorded neurons as the fraction of stimuli that are correctly decoded from the recorded population activity (see Methods). The contribution to the behavioral accuracy in stimulus discrimination that is attributable to the recorded neurons was estimated from our model as the total task performance predicted by our model minus that attributable to neurons that were not recorded. The latter was computed as the behavioral discrimination performance predicted by our readout model after shuffling the values of the neural predictors across trials (see Methods). In the sound localization PPC dataset, we estimated that the ~50 recorded PPC neurons contributed to increase task performance by ~3.5% (Fig 4c). Stimulus information in PPC activity appeared to be read out for behavior, but not optimally and entirely, given that the same population contributed ~10% decoding performance above chance (Fig 4b, left panel). In the evidence accumulation dataset, the ~350 recorded neurons contributed to increase task performance by ~25% (Fig 4e), corresponding again to a significant but not optimal and entire readout of the ~30% stimulus decoding accuracy above chance contributed by the same population (Fig 4d, left panel). Similar considerations would apply for the AC data in the sound localization task (see Extended Data Figure 6,7). Thus, this set of results suggest that the behavioral readout is suboptimal, because it does not use all stimulus information to inform the behavioral choice.

A second possible operational way to define an optimal behavioral readout is to define it as the one in which the behavioral choice in each trial is equal to the stimulus decoded from neural activity. Verifying this hypothesis requires computing how tight is the relationship between the stimulus decoded from neural activity and the behavioral choice. In our data, we computed the readout efficacy, defined as the probability that, in a given trial, the behavioral choice matches the stimulus decoded from population activity (see Methods). According to the above definition, an optimal readout of all stimulus information would have a value of readout efficacy equal to 100%. In our PPC data, the readout efficacy, or probability that the choice matches the decoded stimulus, was 61.0% ± 0.2% in the sound localization dataset and 91.1% ± 0.1% in the evidence accumulation dataset. This empirical observation is compatible with the view that stimulus information in neural activity is read out, but not optimally and entirely, for task performance. It also implies that it is not meaningful to compare the task performance predicted by the enhanced by consistency readout to the one predicted by an optimal readout of stimulus information, defined as a readout in which all choices follow the decoded stimulus. This is why in Figure 4j-n we compared the enhanced-by-consistency readout with the consistency-independent one that, like the optimal readout, is based only on the decoded stimulus, but is equalized for readout efficacy (see Methods, Section "Matching enhanced-by-consistency and consistency-independent readouts in terms of efficacy").

Another way to investigate whether our data are compatible with the hypothesis that the behavioral readout of the recorded population is suboptimal considers the relationship between stimulus and choice information in the same population. In both PPC datasets we found that the recorded PPC neuronal population had more choice than stimulus information (again, we quantified stimulus (choice) information as the fraction correct of a decoder of stimulus (choice) identity). Specifically, in the sound localization dataset, stimulus decoding was 60.1% ± 0.2% correct (Fig. 4b, left) and choice decoding 67.7% ± 0.2% correct (see also Ref[6] for comparable information theoretic results based on single cell analysis). In the evidence accumulation dataset, stimulus decoding was 80.6% ± 0.1% correct (Fig. 4d, left) and choice decoding 96.9% ± 0.1% correct. This suggests that choice is determined, at least in part, by stimulus-unrelated components of the activity of the recorded neural population. This in turn suggests that the readout of the activity of that population is suboptimal in terms of reading out stimulus information, because choice is based also on non- stimulus informative features of population activity.

Finally, if the readout of the sensory information carried by the recorded population was optimal, we would expect that mouse task performance increases monotonically with the posterior probability of the stimulus given the neural activity observed in single trials. Our data instead show (Extended Data Figure 4c-e, g-i) some conflicting evidence: first, that task performance for a given posterior level is not homogeneous across trials and shows a systematic dependence on consistency; second, that the above-mentioned relationship between task performance and posterior probability can be reversed (i.e. higher task performance corresponding to lower values of posterior) for selected subsets of trials (for example, task performance in correctly decoded consistent trials with medium posterior is higher than task performance in correctly decoded inconsistent trials with high posterior, see Extended Data Figure 4d-e, h-i). Both findings are not compatible with the hypothesis of optimal readout.

All in all, our results are all compatible with the notion that the behavioral readout does not use optimally all the information carried by neural activity for task performance.

### Supplementary Information References


1. Pola, G., Thiele, A., Hoffmann, K. P. & Panzeri, S. An exact method to quantify the information transmitted by different mechanisms of correlational coding. *Netw. Comput. Neural Syst.* **14**, 35–60 (2003).
2. Panzeri, S., Schultz, S. R., Treves, A. & Rolls, E. T. Correlations and the encoding of information in the nervous system. *Proc. R. Soc. Lond. B* **266**, 1001–1012 (1999).
3. Averbeck, B. B., Latham, P. E. & Pouget, A. Neural correlations, population coding and computation. *Nature Reviews Neuroscience* **7**, 358–366 (2006).
4. Nogueira, R. *et al.* The effects of population tuning and trial-by-trial variability on information encoding and behavior. *J. Neurosci.* **40**, 1066–1083 (2020).
5. Runyan, C. A., Piasini, E., Panzeri, S. & Harvey, C. D. Distinct timescales of population coding across cortex. (2017). doi:10.1038/nature23020
6. Pica, G. *et al.* Quantifying how much sensory information in a neural code is relevant for behavior. in *Advances in Neural Information Processing Systems 30* (eds. Guyon, I. et al.) 3686–3696 (Curran Associates, Inc., 2017).
